# Addressing anemia severity in antimony-resistant *Leishmania donovani* infection at the nexus of oxidative outburst and iron transit

**DOI:** 10.1101/2024.03.04.583250

**Authors:** Souradeepa Ghosh, Krishna Vamshi Chigicherla, Shirin Dasgupta, Yasuyuki Goto, Budhaditya Mukherjee

## Abstract

Despite the withdrawal of pentavalent-antimonials in treating Visceral leishmaniasis for more than a decade, recent clinical isolates of *Leishmania donovani* (LD) exhibit unresponsiveness towards pentavalent-antimony (LD-R). This antimony-unresponsiveness points towards a genetic adaptation that underpins LD-R’s evolutionary persistence and superiority over sensitive counterparts. This study highlights LD’s response to antimony exposure in terms of increased potential of scavenging host-derived iron within its parasitophorous vacuoles (PV). LD-R employs a strategy to both produce and rapidly scavenge host-iron in a ROS-dependant manner, and selectively reshuffle iron exporter, Ferroportin, around its PV. Higher iron utilization leads to subsequent iron-insufficiency, compensated by increased erythrophagocytosis facilitated by the breakdown of SIRPα, orchestrated by a complex interplay of two proteases, Furin and ADAM10. Understanding these mechanisms is crucial for managing LD-R infections and their associated complications, like anemia, and may also provide valuable insights into understanding resistance developed in other pathogens that rely on host iron.

## Introduction

*Leishmania donovani* (LD), is an intracellular protozoan parasite responsible for causing visceral leishmaniasis (VL), a potentially fatal disease. The traditional treatment for VL, pentavalent antimonials (SbV), has been discontinued in the Indian sub-population for over a decade due to resistance development. Despite this, recent clinical isolates still show unresponsiveness to SbV, indicating ongoing selection pressures (1). Studies have demonstrated that patients infected with antimony-resistant LD (LD-R) tend to have a higher parasite load in their spleen and liver compared to those infected with sensitive strains (LD-S) (2). This suggests that LD-R isolates possess genetic adaptations that confer a survival advantage in the face of drug exposure. Successful LD-R evolution should involve evading host defenses and the anti-leishmanial effects of antimony, allowing it to prevail as persistent strain and proliferate rapidly as intracellular amastigotes within macrophages (MФs) of the spleen/liver & bone marrow of mammalian host as compare to LD-S. As the initial host defense strategy as well as the anti-leishmanial mode of action of antimony hinges on inducing oxidative stress to eradicate LD parasites, it is crucial to consider reactive oxygen species (ROS) as a significant contributor in the selective survival of intracellular LD-R. ROS function as a double-edged sword in shaping the outcome of LD infection: on one hand, they play a vital role in completely eradicating the pathogen, while on the other hand, a controlled level of ROS is necessary for facilitating amastigote differentiation (3). The primary source of reactive oxygen species (ROS) is the multi-subunit complex NADPH oxidase (NOX). This complex consists of six subunits, three of which (p47, p40, and p67) get phosphorylated to become activated and assemble around the phagolysosomal membrane harboring intracellular LD to generate ROS and kill it (4). To date, most of the intracellular pathogens, including *Leishmania*, have been reported to employ strategies to delay or impair NOX assembly and evade oxidative stress (5–7). Therefore, it is appealing to investigate whether LD-R also delays ROS production or employs some unique strategy to combat ROS for its rapid intracellular proliferation. Interestingly, high ROS has been intricately linked to iron, an essential nutrient derived from heme, and required for the proliferation and pathogenicity of *Leishmania* (8). However, Kinetoplastida, including *Leishmania*, have lost the ability to synthesize heme. Although, *Leishmania*, has partially regained the genes for the final three enzymes of the heme biosynthesis pathway through horizontal gene transfer (9), primarily they are heme auxotrophs reliant on host iron scavenging for survival. Thus, it can be hypothesized that LD-R should scavenge host-derived iron within their parasitophorous vacuole (PV) more efficiently than their sensitive counterparts to support essential cellular processes like DNA replication, oxygen transport, and regulation of antioxidant defense resulting in their aggravated proliferation. However, little is known about the molecular events that might contribute to increased iron acquisition while preventing iron efflux from LD-R infected MФs. Thus, to understand the mechanisms underlying iron acquisition and utilization in LD-R infection, a range of iron transporter and metabolizing proteins has been examined that may facilitate its rapid proliferation outcompeting LD-S. Importantly, higher acquisition of host-derived iron can also explain cases of severe anemia, with a more frequent incidence of Coombs-positive hemolytic anemia among VL-positive cases from the Indian subcontinent (10). Furthermore, macrophages also play a key role in iron production by recycling senescent red blood cells (RBCs) through erythrophagocytosis. The fate of erythrophagocytosis is regulated by the interaction between CD47, enriched in live RBCs, and SIRPα, expressed on macrophage membranes, and previously reported to be regulated by replicating intracellular LD through post-translational modifications (11). The mechanism of anemia in VL remains complex, multifactorial, and largely unknown (12), enhanced erythrophagocytosis of both live and dead RBCs may explain severe anemia observed in clinical drug-resistant VL infection. This study aims to identify the potential cause and underpinning mechanism of severe anemia, particularly in VL patients infected with LD-R, both *in vivo* and *in vitro*. Understanding these molecular events could elucidate the mechanism of the rapid emergence of drug resistance not only in VL but also in other intracellular heme-auxotrophic pathogens exploiting similar mechanisms.

## MATERIALS AND METHODS

### I. Reagents and antibodies

Lipofectamine 2000 Reagent (11668019) was purchased from Invitrogen. Giemsa stain solution (72402) was purchased from SRL. DAPI (TC229) was purchased from Himedia. Fluoromount-G (00-4958-02) was obtained from Invitrogen. Harris hematoxylin and eosin were obtained from Merck. SSG is a kind gift from Albert David, Kolkata. Lysotracker Red (L7528) and Calcein-AM (C3099) were purchased from Invitrogen. Antibodies against Rab5a (E6N8S), NRF2 (D1Z9C), Histone (H3) (D1H2), p50 (D4P4D), c-Rel (D4Y6M), FTH1 (D1D4), CD11b (E6E1M), GAPDH (D16H11) and β-actin were purchased from Cell Signaling Technologies (CST). Anti-Mouse CD11b-Alexa Fluor 488 is purchased from Invitrogen (53-0112-82). Antibodies against p47phox (sc-17844) and p50 (sc-8414) were purchased from Santa Cruz Biotechnology. Antibodies against p62/SQSTM1 (MA5-27800) and Hmox1 (PA5-88074) were purchased from Invitrogen. Antibodies against CD71/ Transferrin Receptor (A5865) and Rab5a (A1180) were purchased from Abclonal. Antibody against NRAMP1 was purchased from Santa Cruz Biotechnology (sc-398036). Antibody against Furin was purchased from Abclonal (A5043) and CST (E1W40). Antibody against ADAM10 was purchased from Abclonal (A10438) and Invitrogen (MA5-23867). Anti-Mouse CD172a (SIRP alpha)-APC was purchased from Invitrogen (17-1721-82). Ferroportin/ SLC40A1 antibody was purchased from Novus Biologicals (NBP2-45356). HRP-conjugated anti-mouse secondary antibody (57404) was purchased from SRL and anti-rabbit secondary antibodies were purchased from Sigma (A8275). Alexa Fluor 488 goat anti-mouse (A11001), Alexa-fluor 594 goat anti-mouse (A11005), Alexa-fluor 488 goat anti-rabbit (A11008), and Alexa-fluor 594 goat anti-rabbit (A11037) secondary antibodies were purchased from Invitrogen. Anti-rat IgG (H+L) Alexa Fluor 647 antibody (4418S) was purchased from CST.

### II. Mice and infection

BALB/c mice (*Mus musculus*) were maintained and bred under pathogen-free conditions. The use of mice was approved by the Institutional Animal Ethics Committees of the Indian Institute of Technology, Kharagpur, India. All animal experiments were performed according to the National Regulatory Guidelines issued by biosafety clearance number IE-01/BM-SMST/1.23. a. Metacyclic promastigotes (1×10^7^) were pelleted and dissolved in 200µl sterile PBS and intravenous infection was performed in the tail vein of 4-6 weeks old BALB/c mice using an insulin syringe. Mice were sacrificed at 28 days pi or 24 weeks pi (for establishing the anemia model) as per experimental requirements. Before sacrificing, blood was drawn from the cardiac puncture under anesthesia with isoflurane, and collected in an EDTA-vacutainer. These samples were run in Erba H560 Fully Automated 5-part Hematology Analyser and the hematological parameters were noted. The mice were then sacrificed by cervical dislocation and the spleen was separated for further analysis of the splenic parasite burden and erythrophagocytosis monitoring.

### III. Macrophage (MФ) isolation

Mouse peritoneal MФs were harvested from BALB/c mice by lavage, 48hours (hrs) after intraperitoneal injection of 3% soluble starch (SRL). The adherent peritoneal exudate cells (PECs) were defined as MΦs for convenience. MФs were plated on sterile 12mm round coverslips (Bluestar) in 24-well plate or 35mm dish (Tarsons) or 6-well plate at a density of 1×10^5^ cells/coverslip or 1×10^6^ cells respectively in RPMI 1640 medium (GIBCO) supplemented with 10% FBS (GIBCO) and 100 U PenStrep (GIBCO) (RPMI complete medium) as and when required. The cells were left to adhere for 48hrs at 37°C and 5% CO_2_ before infection.

### IV. Parasites maintenance and infection

*Leishmania donovani* clinical isolates used in this study were as follows: MHOM/IN/83/AG83, MHOM/IN/09/BHU777/0, MHOM/IN/10/BHU814/1, MHOM/IN/09/BHU575/0 (kind gift from Dr.Syamal Roy, CSIR-IICB), and MHOM/NP/03/D10. DD8 is primarily an antimony-sensitive strain while the Amphotericin-B resistant DD8 strain is laboratory-generated and is a kind gift from Dr. Arun Kumar Halder, CDRI, Lucknow, India. D10 (13) is obtained from the Laboratory of Molecular Immunology, Graduate School of Agricultural and Life Sciences, The University of Tokyo, Bunkyo-ku, Tokyo, Japan. Transgenic LD-S GFP and LD-R GFP lines were generated in the lab. All the isolates were maintained in BALB/c mice. Promastigotes were maintained in a 22°C auto shaker incubator in M199 medium supplemented with 10% FBS. Peritoneal MФs from BALB/c mice were infected with LD strains at a ratio of 1:10 for 4hrs infections followed by washing and incubating for a specific time according to experimental requirements and stained with Giemsa.

### V. Parasitophorous vacuole (PV) isolation

PV was isolated following the method described previously (14). Briefly, infected cells were gently scraped off and resuspended in homogenization buffer (20mM HEPES, 0.5mM EGTA, 0.25M sucrose, and 0.1% gelatin supplemented with protease inhibitor cocktail). Lysis was performed by repeated passage (5-20 times) through two 1ml syringes with 23G needles. The mixture was then centrifuged at 200x*g* for 10minutes (mins) to pellet intact cells and nucleus. The low-speed supernatant was then loaded onto a discontinuous sucrose gradient consisting of 3ml 60% sucrose, 3ml 40% sucrose, and 3ml 20% sucrose in HEPES-buffered saline (30 mM HEPES, 100 mM NaCl,0.5 mM CaCl, 0.5 mM MgCl_2_, pH 7) in 15ml falcon tube and centrifuged at 700x*g* for 25mins at 4°C. The parasitophorous vacuoles (PV) were harvested from the 40-60% interface carefully. It was then centrifuged and pelleted at 12,000x*g* for 25mins. The supernatant was discarded and the pellet fraction is PV.

### VI. Measurement of intracellular ROS production

Peritoneal MФs were plated in a 96-well plate at a density of 20000 cells/well and allowed to adhere for 48hrs. MФs were infected with metacyclic promastigotes at a ratio of 1:10 (macrophage: parasite) for 1-8hrs. MФs were loaded with H_2_DCFDA (2’,7’-Dichlorodihydrofluorescein diacetate) at 10µM final concentration and kept for 45mins at 37°C 5% CO_2_ followed by washing in HBSS. DCF fluorescence was considered as a read-out of intracellular ROS. Area-scan readings were taken using a Biotek Cytation 5 imaging reader at Ex/Em:492–495/517–527nm. Each experimental sets were performed in triplicates.

### VII. Quantification of NADPH

NADPH is first extracted from pelleted LD-R and LD-S promastigotes using a method as described elsewhere (15). Briefly Solvent A (40:40:20 acetonitrile: methanol: water supplemented with 0.1M formic acid) was added to the parasite pellet, vortexed for 10seconds (secs), and allowed to rest on ice for 3mins. For each 100 µl of solvent A, 8.7 µl of solvent B (15% NH_4_HCO_3_ in water (w:v) precooled on ice) was then added and vortexed to neutralize the sample. The mixture was kept in dry ice for 20mins. Samples were then centrifuged at 16,000x*g* for 15mins at 4°C and supernatant was taken for LC-MS analysis.

### VIII. Metabolomics Analysis

Mid-log phase promastigotes were used, and metabolite extraction was adapted using the protocol described in t’Kindt *et al* (16). The RPLC-MS was performed using a C18 column as the stationary phase and a methanol-water mixture with increasing concentration over time as the mobile phase. The MS1 spectral data were generated in dual ion mode using Electron Spray ionization. Generated Water Raw files were converted to mzXML files, an open mass spectrometry data format, using the msconvert tool from ProteoWizard (17) filtering only positive ions. A GUI-based Mass Spectrometry Data Processing Engine called El-Maven v0.12.0 (18) was used for subsequent analysis. Peak alignment was performed using the OBI-Warp alignment algorithm and Savitzky–Golay filter for ESI smoothening. The peak annotation was done using the estimated m/z of the selected compounds by their compound formula with a 100-ppm error and 5 best peaks for each group were selected. From the result annotation, the sum of the intensities was used for each metabolite and any peak with a high RT difference was manually excluded for the analysis.

### IX. Gene-Set Enrichment Analysis (GSEA)

GSEA was performed leveraging publicly available Microarray data from NCBI GEO data set: GSE144659 to differentially compare clinical LD-R isolates (BHU575, BHU814) and LD-S isolates (AG83, BHU777). A Group-Wise Mean Normalisation is performed to make the data comparable for all the samples. A Multi-Dimensional Scaling is performed to generate a PCA plot. Using the generated normalized data, and merging it with Glycolysis and Pentose phosphate pathway Gene sets obtained from KEGG, the GSEA analysis is performed using the GSEA GUI software (19, 20).

### X. RNAseq analysis

Differential Expression Analysis (DGE) was performed for LDS 4hrs, LDS 24hrs, LDR 4hrs, and LDR 24hrs post-infected MФs keeping uninfected (UI) MФ as control, using the RNA-Sequencing read count data with DESeq2 library on R (21). RNA-Sequencing was performed utilizing external service by Bionivid Project. The results were represented as a volcano plot by using a cutoff of log_2_(Fold-Change)≥0.6 for upregulated genes and log_2_(Fold-Change)≤-0.6 for downregulated genes (22, 23), and with Benjamini-Hochberg adjusted p-value<0.1. Normalized counts were obtained from the DESeq2 package to develop heatmaps of a few selected genes using gplots and RColorBrewer.

### XI. Calcein-AM-based estimation of Labile iron pool (LIP)

Labile iron pool status was estimated using Calcein-AM as described previously (24, 25). Briefly, 2×10^5^ peritoneal MФs were plated in Confocal dishes for 48hrs followed by infection with LD promastigotes at a specific time point. MФs were then loaded with 0.5µM working concentration of Calcein-AM (Ex/Em:494/517nm) for 30mins at 37°C. It was washed properly with 1XPBS after which MФs were exposed to 50nM Lysotracker Red (Ex/Em:577/590nm) to demarcate the parasitophorous vacuole for 1hr at 37°C followed by washing in 1XPBS. MФs were then incubated with DAPI in 1XPBS followed by washing and taken for Confocal microscopy for live cell imaging.

### XII. Quantification of cytoplasmic and intraphagosomal iron

Iron concentration was quantified by colorimetric ferrozine-based assays as described previously (26). For whole intracellular quantification of iron, infected cells were scraped off. Phagosomes of infected cells were isolated for intra-phagosomal iron quantification and cytoplasmic fractions were isolated for cytoplasmic iron quantification as mentioned earlier. Briefly, cytoplasm/ phagosomes were lysed by application of 200µl 50mM NaOH + 200µl 10mM HCl + 200µl iron releasing reagent (a freshly mixed solution of equal volumes of 1.4M HCl and 4.5% (w/v) KmnO_4_ in H_2_O). The mixtures were kept in the dark and incubated for 2hr at 60°C within a fume hood. 60µl of iron detection reagent (6.5mM ferrozine, 6.5mM neocuproine, 2.5M ammonium acetate, and 1M ascorbic acid dissolved in water) was added after the samples were cooled down and incubated for another 30mins at room temperature. 280µl of solution was added to a 96-well plate and absorbance was measured at 550nm. Iron concentrations were determined using known FeCl_3_ standards. Each experimental sets were performed in triplicates.

### XIII. Plasmids, transfection, and luciferase reporter assay

Using the Eukaryotic Promoter Database (EPD) heme oxygenase-1 promoters were determined. Using the PROMO tool, the transcription factor binding sites were determined for p50 and cRel. Murine HO-1 promoters (−1385/+137), 1522bp using primers 5’-AAGGTACCTGAGGCTGGAGAGATGGCC-3’ and 3’-TAAAAGCTTCACCGGACTGGGCTAGTTCAG-5’ were PCR amplified and cloned in promoterless PGL3 enhancer empty vector (Promega, E1771) at the upstream of luciferase gene. The whole HO-1 promoter was separated into two halves: Site A^−/−^ (−4/−635) and Site B^−/−^ (−636/−1377). Using NEBase Changer tool, two truncated promoter primers (Site A^−/−^ and Site B^−/−^) were designed: Site A−/− (5’-TCAGATTCCCCACCTGTA-3’ and 3’-GGTACCTTTATCGATAGAGAAATG-5’) and Site B−/− (5’-GCTCACGGTCTCCAGTCG-3’ and 3’-GCTGGAGGTTGAAGTGTTC-5’). Two sites were mutated following the Q5 Site-Directed Mutagenesis protocol. RAW 264.7 macrophage cell lines were transiently transfected with HO1+PGL3, SiteA^−/−^ HO1+PGL3, and SiteB^−/−^HO1+PGL3 plasmids using Lipofectamine 2000 reagent (Invitrogen) for 6hrs, washed and incubated for another 12hrs followed by infection with LD-S or LD-R or kept uninfected. Luciferase activity was measured in cell extracts using a Dual-Luciferase Reporter Assay Kit (Promega).

### XIV. Western Blot analysis

Following infection and other treatments, peritoneal MФs were scraped off in ice-cold PBS followed by centrifugation and lysed in 2X SDS sample loading buffer consisting of 20% glycerol, 10% β-mercaptoethanol, 4% SDS, 0.125M Tris HCl, 0.004% Bromophenol blue, pH=6.8. Samples were boiled at 95°C for 10mins and subjected to SDS-PAGE. Resolved proteins were transferred in a nitrocellulose membrane (Bio-Rad) using a Semi-dry transfer apparatus (Invitrogen). Membranes were blocked in blocking solution (5% milk in 0.05%Tween20/PBS) for 30mins at room temperature followed by probing in primary antibodies diluted in blocking solution overnight at 4°C in an orbital shaker. Membranes were then incubated with the following HRP conjugated goat anti-rabbit IgG or goat anti-mouse IgG secondary antibody. The substrate working solution was using SuperSignal West Pico PLUS Chemiluminescent Substrate (Thermo Scientific). The membranes were incubated with the substrate working solution for developing and the chemiluminescence blot images were taken in Chemidoc MP Imaging System (Bio-Rad). GAPDH or β-actin and Histone (H3) were used as a positive control for cytoplasmic fraction or whole cell lysate and nuclear fraction respectively.

### XV. Immunofluorescence

For immunofluorescence studies, infected, uninfected, or treated MФs plated in glass-coverslips were fixed in 2% paraformaldehyde for 5-10mins. The fixative was aspirated and neutralized with 0.1M glycine/PBS for 5mins followed by permeabilization in 0.2% Triton in 1XPBS for 20mins on an orbital shaker. Subsequently, MФs were blocked with blocking solution (2% BSA in 0.2% Triton in 1XPBS) for 20mins on an orbital shaker at 4°C. MФs were then incubated in primary antibodies diluted in blocking solution against specific proteins for 1hr at room temperature on an orbital shaker. MФs were then washed thoroughly for 3 × 5mins with 0.1% Triton/PBS. For differential permeabilization assays, permeabilization-free (no detergent) labeling was performed for membrane protein (in this case CD11b and ADAM10) followed by permeabilization (0.1% Triton in 1XPBS) and labeling with antibodies against the intracellular protein (in this case Furin). Finally, Alexa-Fluor labeled secondary antibodies diluted in blocking solution were loaded onto the MФs and incubated for 1hr at room temperature on an orbital shaker followed by washing 3 × 5mins with 0.1% Triton/PBS. The coverslips were then mounted using Fluoromount+DAPI in glass slides and viewed in a Confocal microscope using an oil immersion 63X objective.

### XVI. *In-vitro* monitoring of erythrophagocytosis

Erythrophagocytosis was monitored using a method used elsewhere (27). Briefly, GFP-tagged LD promastigotes were used for infection on peritoneal MФs (1:10 macrophage: parasite) seeded on 16-well chamber slide glass for specific time points (4hrs or 24hrs). MФs were then incubated with freshly prepared 100µl Cyto-Red RBC and kept for 4hrs at 37°C, 5% CO_2_ followed by being washed and monitored by live cell imaging.

### XVII. RNA isolation from spleen and amastin quantification by qRT-PCR

~100mg of splenic sample were homogenized using a micro-pestle and resuspended in RNAiso Plus (Takara). RNA was isolated using the manufacturer’s protocol and the quality and quantity were estimated in a NanoDrop Lite spectrophotometer (Thermo Scientific). Using ProtoScript First Strand cDNA Synthesis Kit (New England BioLabs), RNA is denatured and cDNA was synthesized. From the diluted cDNA product, using PowerUp™ SYBR™ Green Master Mix and Forward primer: 5’-GTGCATCGTGTTCATGTTCC-3’; and Reverse primer: 3’-GGGCGGTAGTCGTAATTGTT-5’ were subjected to qRT-PCR for amplifying Amastin in QuantStudio 5 (Applied Biosystems) in triplicates. Finally, the amplification plots were generated and analyzed using QuantStudio^TM^ Design & Analysis Software v1.5.2 (Applied Biosystems). The ΔCt and ΔΔCt for Amastin were calculated with respect to Murine β-actin (Forward primer: 5’-AGAGGGAAATCGTGCGTGAC-3’, and Reverse primer: 3’-CAATAGTGATGACCTGGCCGT-5’).

### XVIII. Flow cytometry

Metacyclic promastigotes were sorted using Beckman Coulter Cytoflex srt by gating *FSC-A* × *SSC-A* where FSC^low^ represents the metacyclic population and FSC^high^ represents the procyclic population (28). While GFP-positive P1 populations were enumerated in the Flow cytometer (BD Biosciences). Briefly, 5ml P1 promastigote culture differentiated from spleen-macerates was resuspended in 1ml FACS buffer and run in FACS and the data is recorded in logarithmic scale. For the SIRPα^+^ population enumeration in infected MФs, 1×10^6^ cells were stained by labeling with Anti-Mouse CD172a (SIRP alpha)-APC and Anti-Mouse CD11b-Alexa Fluor 488 followed by washing and fixing with 2% PFA. The cells were resuspended in 500µl ice-cold FACS buffer and run in the Flow cytometer. All analyses were performed in FlowJo v10.

### XIX. Histology staining

Spleen collected at the time of sacrificing the BALB/c mice were fixed with 20% buffered formalin overnight and a representative section was subjected to tissue processing; embedded in paraffin the next day. Once the paraffin mold was completely dried, the tissue block was cut at 4µm thickness in Microm HM 315 Microtome and placed in Mayer’s egg albumin-positive slides. Once fully dried, the slides were deparaffinized using xylene followed by dehydration, and then stained with hematoxylin for 30secs. It was then rinsed with running tap water for 30mins followed by eosin staining for 10secs. It was rinsed again under running tap water for 20mins followed by dehydration and mounting in D.P.X (Sigma) and observed under the microscope.

## Results

### 1. Antimony-resistant *Leishmania donovani* (LD-R) outcompete drug-sensitive isolates (LD-S) in an experimental model of murine infection

Earlier studies reported that infection with antimony-resistant clinical LD isolates (LD-R) always leads to higher organ parasite load in mice, a condition also reflected in human VL patients (2, 29). Formerly, increased metacyclogenesis observed in LD-R was reported as the sole known contributor in terms of higher *in-vivo* infectivity potential resulting in higher organ-parasite burden (2, 28, 29). However, it has been reported that LD-R promastigotes do exhibit higher replicative potential as compared to their sensitive counterparts (LD-S) (30), which might also provide them with a selective intracellular survival advantage resulting in higher organ parasite load. Thus, to check this, an equal number of metacyclic promastigotes were sorted for GFP-expressing-LD-S (AG83) and RFP-expressing-LD-R (BHU 575) (**Figure 1A.i. and A.ii**.) using Beckman Coulter Cytoflex srt and *in-vitro* competitive infection was performed in peritoneal murine MФs by giving a mixed infection of LD-S GFP and LD-R RFP (**Figure 1A.iii**.) to determine their capability to invade and colonize host MФs. Infected MФs were washed after 4hrs post-infection (pi), and kept for another 24hrs. Live cell imaging and videography were performed to record the initial entry and further colonization of LD parasites by checking GFP and RFP-positive amastigotes (**Figure 1A.iv., Supplementary Video 1**). While an equal number of LD-S GFP and LD-R RFP promastigotes invade host MФs at 4hrs pi, at 24hrs pi, a significant increase in intracellular LD-R RFP amastigote population was observed suggesting that there might be other factors apart from infectivity contributing towards increased proliferation and intracellular survival of LD-R in the infected host outcompeting their sensitive counterparts. To determine if this increased survival fitness is not restricted to only BHU575, MФs were infected with an equal number of sorted metacyclics for two clinical LD-R (BHU575, BHU814) and LD-S (AG83, BHU777) isolates (**Supplementary Figure 1A**.). Same as before, infected MФs were washed after 4hrs pi to remove loosely attached parasites, and the number of intracellular amastigotes was enumerated for 4hrs and 24hrs pi (**Supplementary Figure 1B.i. and B.iii**.). The result revealed no significant difference between LD-S and LD-R isolates in terms of initial infectivity at 4hrs pi. (**Supplementary Figure 1B.i. and B.ii**.), however, infection with both LD-R isolates resulted in a significantly higher number of intracellular amastigotes 24hrs pi (**Supplementary Figure 1B.i., B.ii., and B.iii**) suggesting this higher intracellular survival in host MФs might be a common attribute of LD-R infection. Since both the LD-R strains reflected a comparable number of intracellular amastigotes, BHU575 was selected as representative LD-R and AG83 as representative LD-S to conduct all further comparisons with the inclusion of other representative strains as and when mentioned. Next, this observation was challenged *in vivo* by performing experimental murine (BALB/c) infection with an equal number of sorted metacyclic LD-R and LD-S promastigotes as described before. Mice were infected either with LD-S GFP (Group 1), with LD-R (Group 2), with a mixture of LD-S GFP: LD-R in an equal 50:50 ratio (Group 3), or with LD-S GFP: LD-R in 80:20 ratio (Group 4). Infected mice (N=3) were sacrificed after 28 days pi and the spleen was isolated, weighed, and cut into 3 sections as represented in (**Figure 1B.i**.) keeping the uninfected spleen as a control. The first section weighing ~ 100mg for each group was used for quantifying *amastin* expression by qRT-PCR to enumerate splenic parasite load (31) (**Figure 1C.i**.). The second section was macerated, stained with DAPI, and observed in fluorescence microscopy to determine the amastigote type and expressed as LDU (LDU= amastigote per nucleated cell * organ weight in mg) (**Figure 1C.ii. and C.iii**.). The third section was macerated and kept in M199 medium with 10% FCS as represented in the scheme (**Figure 1B.i.),** to allow differentiation of LD-amastigotes to promastigotes and the GFP-positive promastigote population from 1^st^ passage was enumerated by flow cytometry. (**Figure 1C.iv**.). Firstly, a significantly larger infected spleen with increased weight was observed in Group 2 (LD-R) mice as compared to Group 1 (LD-S GFP) (**Figure 1B.ii**.). This increase in spleen size and weight for Group 3 and Group 4 infected mice was comparable and again significantly higher than Group 1 mice suggesting that they might have a higher infection load as compared to Group 1. This observation was also supported by relative amastin expression which reflected the highest splenic amastigote load for Group 2 infected mice, followed by Group 3 and Group 4, all of which are significantly higher compared to Group 1 infected mice (**Figure 1C.i**.). Fluorescence microscopy of the infected spleen (**Figure 1C.ii**.) revealed GFP-positive amastigotes around the splenic MФs in Group 1 infected mice whereas in Group 2 there was little or no signal of GFP as anticipated. Both in Group 3 and Group 4 infected spleen there was a significant reduction in the number of GFP-positive amastigotes as expressed quantitatively in LDU (**Figure 1C.iii**.). Furthermore, to confirm the loss of GFP signal in splenic MФs for Group 3 and Group 4 is not due to plasmid loss, another mixed infection of LD-S GFP and LD-S untagged parasites (50:50) was performed in mice same as before (Group 5), which resulted in a significant retention of GFP signal among splenic amastigotes (**Supplementary Figure 1D.i**.) suggesting that this GFP loss in the case of Group 3 and Group 4 is not due to plasmid loss. Finally, flow cytometry-based quantification of the individual group (**Figure 1C.iv.),** revealed ~ 22.7% of the GFP-positive population in the case of Group 1 (LD-S GFP), 0.42% in LD-R (Group 2), which gets significantly reduced to 2.26% for LD-S GFP: LD-R (Group 3, 50:50), 3.55% in Group 4 (LD-S GFP: LD-R 80:20), while for Group 5 GFP positive population was maintained at ~10.3% further confirming the notion that LD-R has an inherent potential to proliferate more surpassing its sensitive counterparts. It is also interesting to note that it took approximately 11-13 days for Group 1, 5-7 days for Group 2, and 7-10 days for Group 3 and Group 4 for transformed LD to appear as promastigotes in culture. SbV susceptibility against independent transformed LD lines for each experimental group showed an EC_50_ ~21.3 and ~20.43 respectively for LD lines emerging from Group 3 and Group 4, similar to the EC_50_ value (~21.6) of the LD-R parental line (**Supplementary Figure 1C and Table 1**). As a whole, these observations thus clearly confirm that higher organ parasite load in case of LD-R infection might be a consequence of their increased intracellular survival in host allowing them to outcompete their sensitive counterparts both *in-vitro* and *in-vivo*.

**Figure 1:**
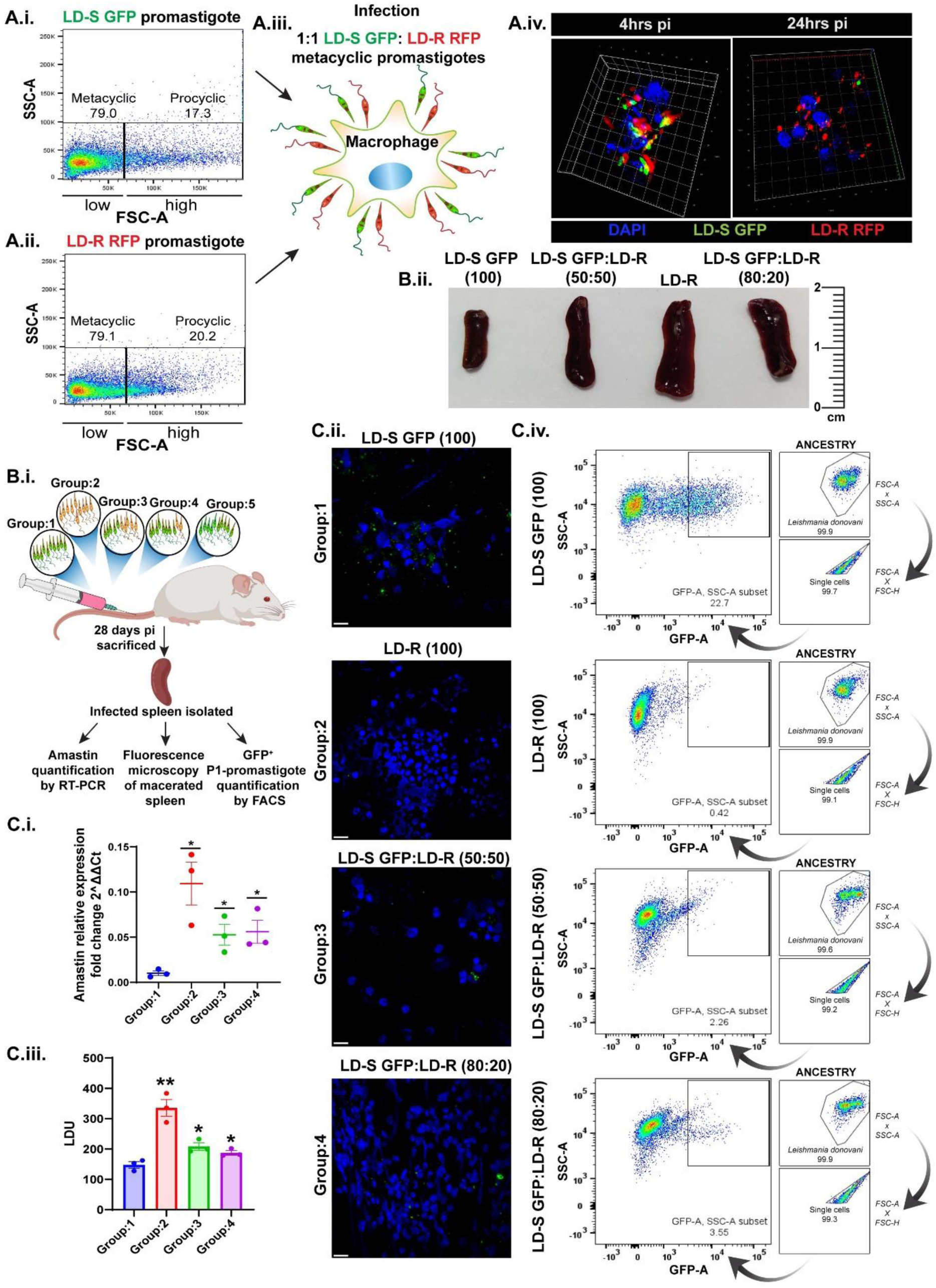
LD-R outcompetes LD-S both in *in-vitro* and *in-vivo* infection: Representative percentage of sorted metacyclics and procyclic LD promastigote population in the culture of (A.i.) GFP-expressing antimony-sensitive (LD-S GFP) and (A.ii.) RFP expressing antimony-resistant (LD-R RFP) sorted in Beckman Coulter cytoflex srt sorter where FSC^low^ (left) represents the metacyclic population and FSC^high^ (right) represents the procyclic population. **(A.iii.)** Scheme representing infection of an equal number of LD-S GFP and LD-R GFP metacyclic promastigotes in peritoneal murine macrophages (MФs). **(A.iv.)** 3-D Z-stack rendering of Super-resolution image showing an equal number of invading LD-S GFP and LD-R RFP parasites in MФs at 4hrs pi (left panel). The right panel shows LD-R RFP amastigote outcompetes LD-S GFP at 24hrs pi. **(B.i.)** Scheme showing tail vein infection of BALB/c mice with LD-S GFP (Group:1), or with LD-R (Group:2), or with LD-S GFP: LD-R in 50:50 (Group:3), or with LD-S GFP: LD-R in 80:20 (Group:4), or with LD-S GFP: LD-S in 50:50 (Group:5). 28 days pi mice were sacrificed and the spleen was isolated from each experimental mouse which was weighed and cut into 3 pieces for performing i. Amastin quantification by qRT-PCR; ii. Fluorescence microscopy of macerated spleen; and iii. GFP-positive population enumeration by flow cytometry. **(B.ii.)** Representative spleen image of infected spleen 28 days pi from mice representing Groups: 1, 3, 2, and 4 (from left to right) with a scale bar (cm) at the right to compare the size of the spleen. **(C.i.)** Relative expression fold change (2^ΔΔCt) of amastin in the infected spleen isolated 28 days pi from Groups:1-4 (N=3 for each group) was determined from qRT-PCR which denotes the amastigote load in each set taking murine β-actin as housekeeping control. **(C.ii.)** Bar graph showing LDU in each experimental set (Groups:1-4) calculated as LDU= amastigote per nucleated cell * organ weight in mg. N=3 for each group **(C.iii.)** Confocal images of the macerated spleen sample 28 days pi for each experimental group (Groups:1-4) representing the GFP amastigote load (small green dots) in each splenic sample. Scale Bar represents 20µm. **(C.iv.)** % of the GFP-positive population was enumerated from flow cytometry to denote the load of LD-S-GFP in comparison to LD-R in experimental Groups 1-4. The right-most panel shows the ancestry of each analysis. Each experiment was performed in 3 biological replicates and graphical data are represented as Mean with SEM. P ≤ 0.01 is marked as **, P ≤ 0.05 is marked as *.

**Table 1:** Table showing EC_50_ value against SbV as determined from the dose-response curve (Supplementary Figure 1C).

| Samples | LD-S GFP | LD-R | LD-S GFP:<br>LD-R (50:50) | LD-S GFP:<br>LD-R (80:20) |
| --- | --- | --- | --- | --- |
| EC50 | 1.787 | 21.6 | 21.3 | 20.43 |
| EC50 range | 1.538 to 2.058 | 20.50 to 22.76 | 20.41 to 22.24 | 19.73 to 21.15 |

### 2. LD-R infection rather than suppressing host-induced reactive oxygen species promotes ROS induction as opposed to LD-S infection

Since both the anti-leishmanial mode of action of pentavalent antimonials as well as the host defense strategy against LD infection primarily involves reactive oxygen species (ROS), we speculated that LD-R might have evolved a mechanism to suppress intracellular ROS generation in infected hosts allowing its higher intracellular survival. To test this possibility, the intracellular ROS was quantified from LD-R and LD-S infected MФs at different time points 2hrs, 4hrs, and 8hrs pi. Remarkably, Confocal microscopy as well as fluorescence plate reader-based quantification revealed an initial high ROS at 2hrs and 4hrs pi in LD-R infected MФs as compared to LD-S infected ones which subsequently flattens out at 8hrs pi (**Figure 2A.i. and A.ii**.). However, incubation of MФs with killed LD-R parasites failed to generate substantial ROS as compared to live LD infection, implying replicating parasites are required for ROS generation. This observation clearly suggests that LD-R as opposed to suppressing intracellular ROS, rather promotes it and still manages to thrive better in this hostile oxidative environment. It is well established that LD-S prevents NOX assembly in the phagolysosomal membrane to delay ROS production (7). Hence, the status of p47 phox, a subunit of NOX was monitored in case of LD-R infection as a readout of NOX activation. A significant co-localization of p47 with Rab5a (early endosomal marker) surrounding LD-R in infected MФs was noticed at 4hrs pi (**Figure 2B**.), which was significantly less in the case of LD-S-PV as reported previously (32). This observation further confirms that LD-R instead of preventing ROS production, elicits ROS production and must have devised a strategy to somehow withstand this initial ROS outburst. (**Figure 2C**). Interestingly, this trait of enhanced early ROS induction seems to be specifically linked with primary antimony unresponsiveness as infection with lab-generated amphotericin-B unresponsive LD isolates (AmpB-R-DD8) without primary unresponsiveness towards antimony failed to elicit ROS production in infected MФs. In contrast, LD-R strain with unresponsiveness only towards antimony (D10) or towards multiple other drugs but with primary unresponsiveness towards antimony could successfully induce ROS production (**Figure 2C**.), suggesting elicitation of ROS outburst followed by neutralization is a trait specific to antimony unresponsive LD infection. To thrive in high ROS, it is anticipated that LD-R must have a strong inbuilt anti-oxidative response. Primary reducing equivalents, NADPH have been widely reported previously to neutralize intracellular ROS by donating reductive potentials to glutathione and thioredoxins (33). As maximum ROS generation was observed in 2-4hrs pi when LD persists as intracellular promastigotes in early PV (5), intrinsic NAPDH level was compared between metacyclic LD-S and LD-R (**Figure 2D, and 2D inset**). HPLC-based quantification of NADPH/ NADP+ in LD-S and LD-R suggests that the NADPH/NADP+ ratio is significantly higher in LD-R as compared to LD-S at a retention time of around 1.5-2mins. To confirm the role of the intrinsic high level of NADPH which might allow LD-R to withstand high ROS, LD-R and LD-S were first exhausted of inherent NADPH by treating with 100µM H_2_O_2_ which has been previously reported to have no effect in LD growth (34), and then incubated with 6-Aminonicotinamide, an inhibitor of G6PD, the rate-limiting enzyme of pentose phosphate pathway shunt (PPP) which produce NADPH. MФs infected with these NADPH-deficient parasites resulted in no significant difference in initial infectivity at 4hrs pi between LD-R, LD-S, NADPH-deficient LD-S, and NADPH-deficient-LD-R, suggesting H_2_O_2_ or 6-AN treatment does not affect their infectivity (**Figure 2E.i. left panel, and E.ii**.). Contrarily, at 24hrs pi, there was a significantly low number of intracellular amastigotes in NADPH-deficient LD-R, even lower than LD-S infection, while for NADPH-deficient LD-S infection, a slight increase in parasite burden was observed in 24hrs as compared to 4hrs pi. (**Figure 2E.i. Right panel, E.ii**.), Measurement of intracellular ROS in MФs infected with NADPH-deficient and untreated LD-R revealed a continuous persistence of intracellular ROS in NADPH-deficient LD-R similar to LD-S even after 8hrs pi (**Figure 2F)** pointing towards the role of intracellular NADPH in neutralizing high levels of persistent ROS in LD-R infected MФs. This observation also suggested that LD-R as compared to LD-S might be preferentially utilizing PPP which acts as the major source of NADPH.

**Figure 2:**
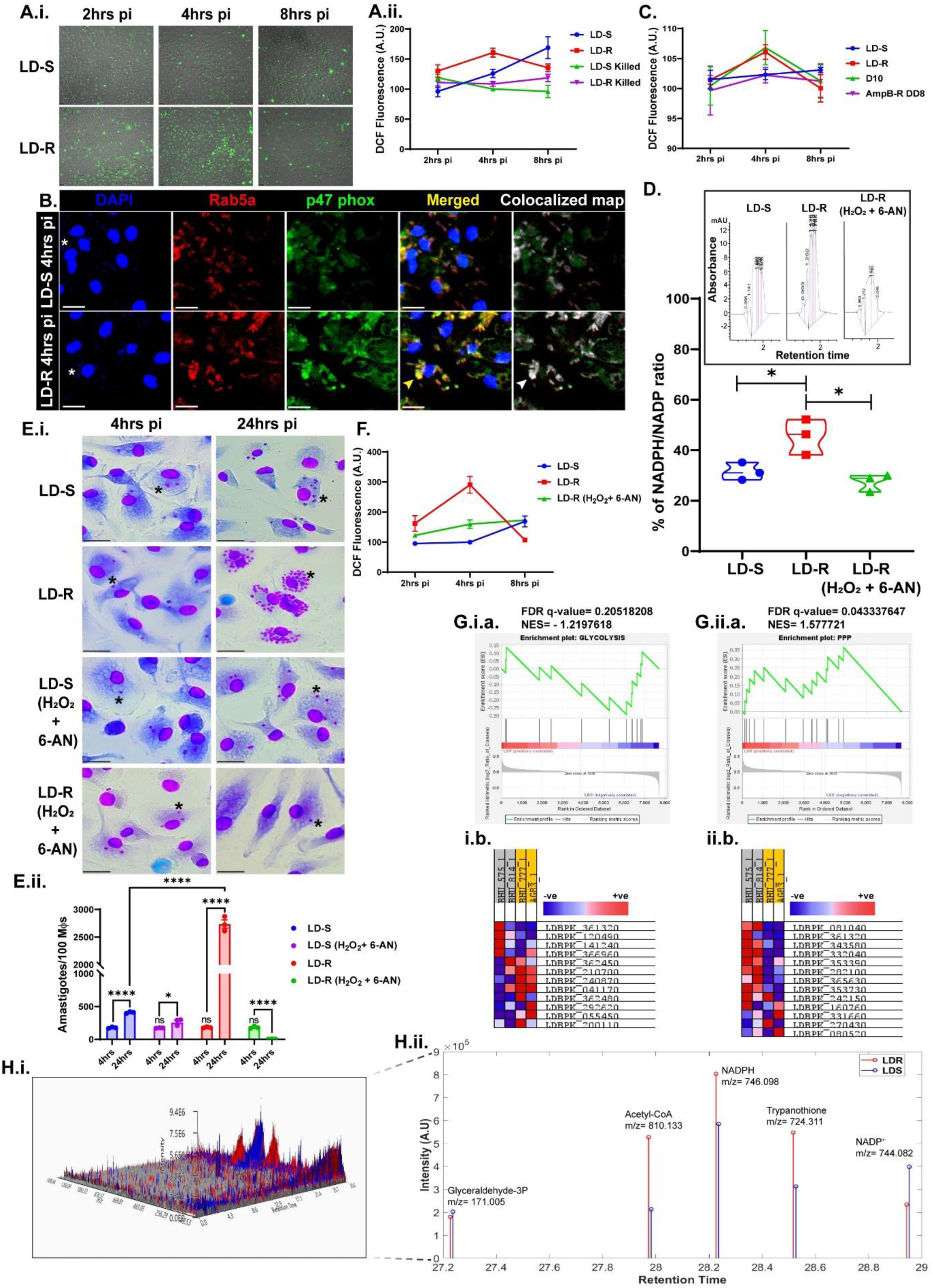
An inherent metabolic shift in LD-R helps these parasites neutralize host-generated ROS outbursts. **(A.i.)** 20X magnification live-cell confocal images of LD-S and LD-R infected MФs at 2, 4, and 8hrs pi image by staining with H_2_DCFDA (green), a read-out of intracellular ROS level merged with DIC. **(A.ii.)** DCF fluorescence quantification of LD-S, LD-R, heat-killed LD-S, and heat-killed LD-R infected MФs at early hours using Biotek Cytation 5 imaging reader. **(B.)** Confocal images showing p47 phox (green) colocalization around early PV demarcated by Rab5a (Red) in LD-S and LD-R infected MФs at 4hrs pi as a read-out of NOX activation. The colocalized pixel map on the right-most panel shows the colocalized region of p47 phox and Rab5a marked in white, the intensity of which is directly proportional to the percentage of colocalization. Yellow Arrow in the 4^th^ panel and white arrow in the 5^th^ panel denote heavily colocalized regions. **(C.)** DCF fluorescence quantification of different strains LD-S, LD-R, D10 (antimony-resistant), and AmpB-R DD8 (Lab generated Amphotericin-B resistant DD8) infected MФs at 2, 4, and 8hrs pi. **(D.)** Graph representing % of NADPH/NADP turnover in LD-S, LD-R, or LD-R (treated with H_2_O_2_ + 6AN to exhaust NADPH) metacyclic promastigotes were quantified in HPLC using NADPH and NADP standards, with inset plot representing the HPLC chromatogram showing the NADPH peak where X-axis denotes retention time and Y-axis denotes absorbance. **(E.i.)** Giemsa-stained images of LD-S, LD-R, NADPH^exh^ LD-S (by H_2_O_2_ + 6AN treatment), and NADPH^exh^ LD-R (by H_2_O_2_ + 6AN treatment) infected MФs at 4hrs and 24hrs. LD nucleus has been marked in (**\***) to show the infected MФs. **(E.ii.)** Amastigotes/100 MФs representing experimental data set as in D.i. were calculated taking different fields. **(F.)** DCF fluorescence quantification of LD-S, LD-R, and NADPH^exh^ LD-R (by H_2_O_2_ + 6AN treatment) infected MФs at early hours using Biotek Cytation 5 imaging reader. **(G.)** GSE analysis showing enrichment plot of glycolysis **(G.i.a.)** and PPP **(G.ii.a.)** in LD-R as compared to LD-S. Glycolysis showed a negative enrichment with a Normalised enrichment score (NES) of – 1.2197618 and FDR q-value of 0.20518208 while PPP showed a positive enrichment with NES of 1.58 and FDR q-value of 0.043337647. **(G.i.b. and G.ii.b.)** Heatmaps showing the expression pattern of the genes in i.glycolysis ii.PPP with • positive enrichment and • negative enrichment of individual strains. **(H.i.)** A 3D snapshot of the whole metabolome showcasing all the resolved and MS data of the run where red denotes LD-R and blue denotes LD-S metabolites. **(H.ii.)** Stem plot showing identified and annotated metabolites having specific m/z values with retention time on the X-axis and their intensity on the Y-axis where red denotes LD-R and blue denotes LD-S metabolites. The scale bar indicates 20µm. One representative small nucleus of LD has been marked in (**\***) to show the infected MФs in Figure 2B and E.i.. Each experiment was performed in 3 biological replicates and graphical data are represented as Mean with SEM. P > 0.05 is marked as ‘ns’ (non-significant), P ≤ 0.05 is marked as *, P ≤ 0.01 is marked as **, and P ≤ 0.001 is marked as ***.

To confirm this hunch, the available microarray dataset, GSE144659, was leveraged and Gene-Set Enrichment Analysis (GSEA) was performed comparing clinical LD-R isolates (BHU575, BHU814) and LD-S isolates (AG83, BHU777) since we could re-confirm previously reported antimony susceptibility for these isolates (35) (**Figure 2G)**. PCA plot was generated for LD-R and LD-S isolates which reflected separate clustering for LD-R (BHU575, BHU814) and LD-S (AG83, BHU777) (**Supplementary Figure 1E)**. Clinical LD isolates BHU138_Clone1 and Clone2, BHU581, and BPK 206 were excluded from this analysis since they were not available to confirm their previously reported susceptibility against antimony. GSE analysis for glycolysis showed a down-regulation in LD-R isolates as compared to LD-S isolates with an enrichment score of – 0.29430503, and a normalized enrichment score of – 1.2197618, with genes LDBPK_200110 (phosphoglycerate kinase C, glycosomal), LDBPK_055450 (pyruvate kinase), LDBPK_292620 (ATP-dependent phosphofructokinase), LDBPK_362480 (glyceraldehyde 3-phosphate dehydrogenase, cytosolic), LDBPK_041170 (fructose-1,6-bisphosphatase, cytosolic, putative), LDBPK_240870 (triosephosphate isomerase) forming the leading edge subset genes for the Glycolysis pathway showcasing effective contribution towards downregulation (**Figure 2G.i.a)** with FDR q-value of 0.20518208. While for PPP, GSE analysis revealed an upregulation for both the LD-R isolates, with an enrichment score of 0.3644994 and a normalised enrichment score of 1.577721, comprising of genes LDBPK_081040 (phosphoribosylpyrophosphate synthetase), LDBPK_361320 (Fructose-bisphosphate aldolase), LDBPK_343580 (phosphomannomutase-like protein), LDBPK_332040 (phosphoribosylpyrophosphate synthetase, putative), LDBPK_353390 (6-phosphogluconate dehydrogenase, decarboxylating), LDBPK_282100 (ribose 5-phosphate isomerase, putative), LDBPK_365630 (phosphoribosylpyrophosphate synthetase, putative), LDBPK_353730 (ribulose-phosphate 3-epimerase, putative), LDBPK_242150 (transketolase), LDBPK_160760 (transaldolase), LDBPK_331660 (ribulose-phosphate 3-epimerase), LDBPK_270430 (Ribokinase, putative), LDBPK_080520 (ribose-phosphate pyrophosphokinase, putative) forming the leading edge subset genes of the PPP pathway showcasing effective contribution towards the upregulation(**Figure 2G.ii.a**) with FDR q-value of 0.043337647. Furthermore, the downregulation of Glycolysis and upregulation of the PPP and pathway in LD-R isolates is also reflected by low FDR values, pointing towards an inherent metabolic shift in LD-R towards PPP. Heatmaps denoting the expression pattern of PPP and Glycolysis pathway genes in all four samples (**Figure 2G.i.b and 2G.ii.b**) show a large cluster of PPP genes positively enriched in LD-R (BHU 575 and BHU 814), and the entire set of genes represented in **Supplementary Table 1A and 1B**.

Taken together these data indicate a significant positive enrichment of PPP in LD-R as compared to LD-S while a negative enrichment of Glycolysis in LD-R as compared to LD-S. To further validate this observation, targeted metabolomics from extracted metabolites of Central carbon metabolism was performed to compare the status of the same in LD-S and LD-R isolates by LC-MS. A 3D plot representing the coverage and total metabolite extraction for both LD-R and LD-S (**Figure 2H.i**.) showed an efficient extraction. On analysis, annotation and quantification of the total metabolomics data for specific metabolites revealed a significant enrichment of NADPH, trypanothione (ROS-neutralizing metabolite), and acetyl CoA, while no significant change was observed for glyceraldehyde-3-phosphate, and a significant decrease in NADP+ level in LD-R as compared to LD-S (**Figure 2H.ii**.). Collectively, both the transcriptomics and metabolomics data incline towards the notion that there must be an inherent metabolic switch evolutionary in clinical LD-R isolates towards PPP which helps them to thrive better in ROS-enriched environments.

### 3. ROS acts as a beneficial tool to produce labile iron in LD-R-infected macrophages

ROS in general has been widely reported to be linked with the killing of intracellular pathogens including LD (36). However, since a significantly higher level of initial ROS in LD-R infected MФs was observed, its role in the higher proliferative potential of LD-R was investigated. LD-S and LD-R infection were performed in MФs in the presence of a suboptimal dose of antimony ~1.52µg/ml (equivalent to EC_50_ value for LD-S(AG83) (35). While this suboptimal dose of antimony resulted in a significant decrease of intracellular amastigotes for LD-S as anticipated, howbeit resulted in a significant increase in the number of intracellular LD-R amastigotes as compared to untreated control 24hrs pi (**Figure 3A.i., top 2^nd^ panel and A.ii**.). However, in presence of N-acetyl-L-cysteine (NAC), an antioxidant, resulted in a significant decrease in the number of intracellular LD-R amastigotes, as opposed to LD-S, which showed a significant increase in the number of intracellular amastigotes in presence of NAC (**Figure 3A.i., 3^rd^ panel and A.ii**.). Similar infection with NADPH^exh^ LD-R by treating with H_2_O_2_+6-AN along with a suboptimal dose of SbV resulted in drastic abolishment of LD-R amastigotes suggesting that inherent NADPH of LD-R helps these parasites to thrive and proliferate in high ROS (**Figure 3A.i., bottom panel and A.ii**.). Taken together, these results imply that ROS might be beneficial for intracellular LD-R proliferation as opposed to LD-S infection since these parasites are shielded with enriched reductive potentials. A previous report on *Trypanosoma* infection in mice also reported a similar observation that oxidative stress aids in parasite persistence (37) by promoting the breakdown of ferritin which stores iron as Fe^3+^ to mobilize bioavailable Fe^2^+ (38). Fe^2+^ is a vital nutrient for LD replication, and LD being a heme auxotroph, relies upon the host for its acquisition. Comparison of Ferritin levels between LD-S and LD-R infected MФs 4hrs pi showed no major change in Ferritin (FTH1) expression (**Figure 3B**), suggesting Fe^2+^ mobilized from Ferritin might not be the source of higher iron to support aggressive replication of LD-R. Apart from Ferritin, another source of Fe^2+^ can be provided by Heme oxygenase-1 (HO-1) which breaks down heme into iron (Fe^2+^) (9, 39). As opposed to Ferritin, a consistently increased HO-1 expression was observed in LD-R as compared to LD-S infected MФs at 4hrs pi (**Figure 3B**). Interestingly, *in silico* analysis with Eukaryotic promoter database and PROMO revealed p50 (−1284/−1277, −635/−624, and −483/−476) and c-Rel (−1230/−1211, −913/−904, −633/−624, and −603/−594) binding site in the promoter region of *hmox-1* gene coding for HO-1(**Figure 3C**.). It has been previously reported that infection with LD-R and not by its sensitive counterpart results in specific activation of p50/c-Rel dependent transcriptional activation in infected MФs (40). Western blot analysis, coupled with confocal microscopy showed a significant co-translocation of p50/c-Rel in the nucleus of LD-R infected MФs as early as 4hrs pi corroborating with the activation of HO-1 (**Figure 3D.i. and D.ii**.). Finally, Luciferase reporter assay with wild type, or truncated promoter constructs: Site A^−/−^ (−4/−635) and Site B^−/−^ (−636/−1377) in the presence of LD-R and LD-S infection, further confirmed specific interaction of p50/c-Rel with Site A of HO-1 promoter resulting in its specific upregulating in LD-R infected MФs (**Figure 3E**).

**Figure 3:**
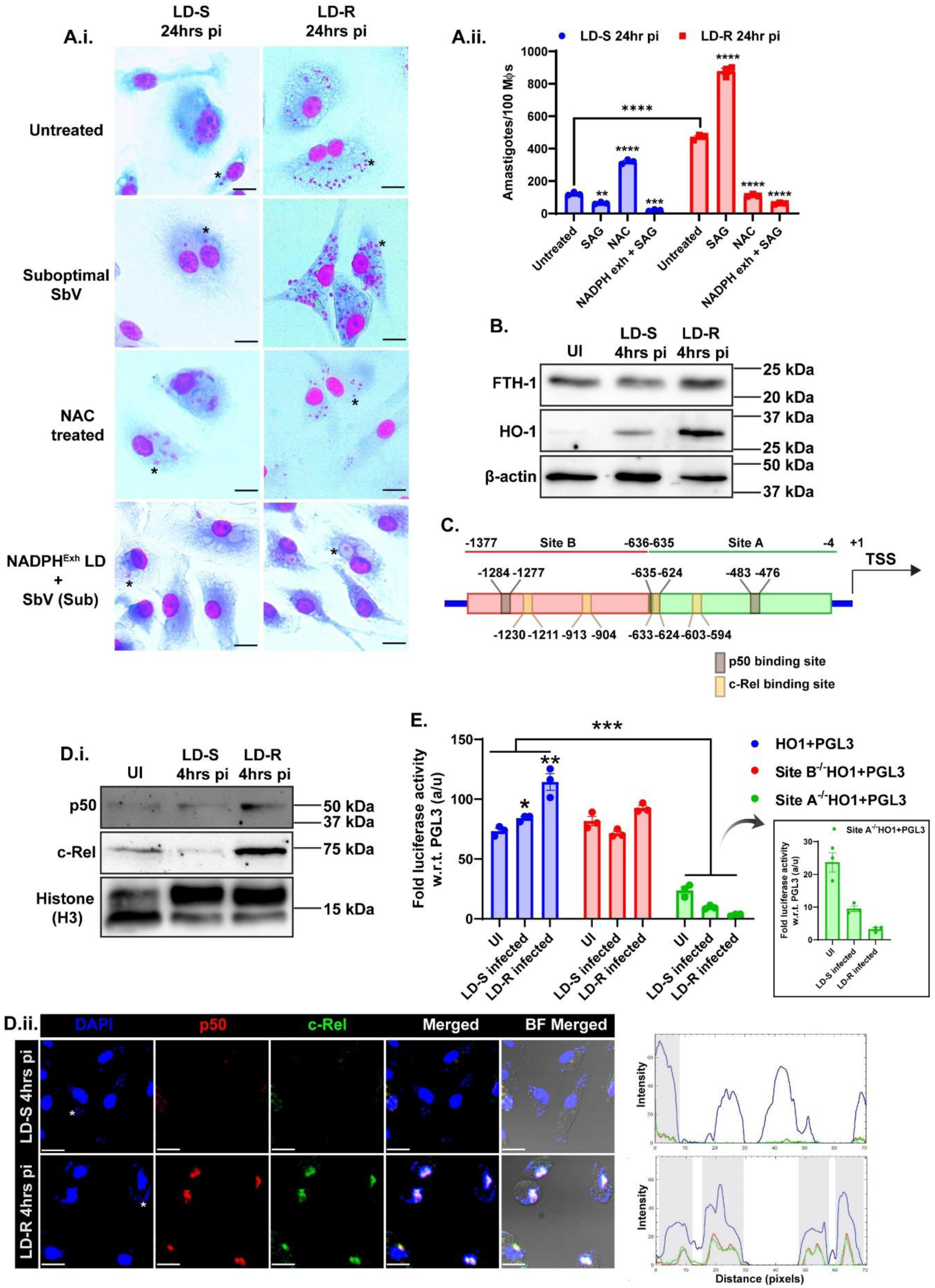
LD-R proliferates more in high ROS by upregulating HO-1 activity. **(A.i.)** Giemsa-stained images of LD-S, LD-R, in murine peritoneal MФs with or without the presence of suboptimal dose of SbV (1.52µg/ml). In some experimental condition LD infection was performed in the presence of N-acetyl-L-cysteine (NAC), while in some experimental conditions, LDs are pre-treated with H_2_O_2_ + 6-AN. All the infections were performed for 24hrs **(A.ii.)** Bar graph showing amastigotes/100MФs in all the experimental sets mentioned in A.i. **(B.)** Western blot of whole cell lysate showing significant high expression of Heme-oxygenase 1 (HO1) in LD-R infected MФs while no significant change in expression level was observed for Ferritin (FTH1) at 4hrs pi. β-actin is used as the positive control. **(C.)** Schematic representation of HO-1 promoter region with p50 and c-Rel binding sites. Site A (−4/−635) was demarcated as green, and Site B (−636/−1377) was demarcated as red. **(D.i.)** Western blot of nuclear fraction showing significant high expression of p50 and c-Rel in LD-R infected MФs 4hrs pi with Histone (H3) as loading control. **(D.ii.)** Confocal images representing the localization of p50 and c-Rel, in MФs, with DAPI representing the nucleus. The right-most panel shows the RGB-profile plot with grey regions demarcating the region where p50 and cRel are colocalized with DAPI. p50 and c-Rel were found to be localized in the nucleus of LD-R-infected MФs. One representative small nucleus of LD has been marked in (**\***) to show the infected MФs. **(E.)** Fold luciferase activity of RAW264.7 cell lysate transfected either with HO1+PGL3 promoter, or Site A (^−/−^), Site B(^−/−^) deleted constructs followed by infection with LD-S and LD-R for 4hrs. PGL3 enhancer empty vector is used for normalization for each transfected set. Each experiment was performed in 3 biological replicates and graphical data are represented as Mean with SEM. P ≤ 0.05 is marked as *, P ≤ 0.01 is marked as **, P ≤ 0.001 is marked as ***, and P ≤ 0.0001 is marked as ****.

### 4. LD-R scavenges labile iron in their PV through reshuffling of host Ferroportin

Given that we observed a significantly higher HO-1 expression in LD-R infected MФs at 4hrs pi, it is anticipated that it should result in an increased labile iron pool (Fe^2+^) in them as compared to infection with their sensitive counterparts. Live cell imaging was performed to measure Calcein-AM fluorescence intensity in LD-S and LD-R infected MФs which acts as a readout of the intracellular labile iron pool (24, 25, 41) (**Figure 4A.i**.) coupled with fluorometric plate-reader-based quantification (**Figure 4A.ii**.)revealed a significantly higher Calcein-fluorescence in LD-R infected MФs as compared to LD-S infection or uninfected control, indicating a significantly diminished labile iron pool in LD-R infected MФs (**Figure 4A.i., ii., and iii**.). This poses an inevitable question if there is an overproduction of iron in response to LD-R infection by HO-1 activation (**Figure 3B**), then why does the labile iron pool get diminished? To track whether Fe^2+^ is infiltrating inside the parasitophorous vacuole (LD-R-PV) where these parasites reside and proliferate, two different approaches were performed: (i) labeling the PV with lysotracker red coupled with Calcein-AM to assess the labile iron pool inside PV by live cell microscopy (**Figure 4B.i**.); (ii) Extraction of PV and cytoplasmic fraction following sucrose density gradient centrifugation coupled with ferrozine-based colorimetric quantification of iron in response to LD-R and LD-S infection (42) (**Supplementary Figure 2A and Figure 4B.ii**.). Live cell imaging revealed early PV harboring LD-R infected MФs showed a significantly low calcein fluorescence, suggesting rapid infiltration of iron takes place in early PV of LD-R infected MФs from cytoplasm as opposed to LD-S infection at 4hrs pi (**Figure 4B.i**.). This observation is also supported by Ferrozine-based quantification of iron which confirmed a significant increase in Fe^2+^ in early PV of LD-R infected MФs with low levels of Fe^2+^ in the cytoplasm as compared to LD-S infected MФs (**Figure 4B.ii**.). Interestingly enough for LD-R infected MФs, a slight increase in Fe^2+^ level in cytoplasm occurs in 24hrs pi along with a significant drop in LD-R-PV as compared to 4hrs pi probably indicating a rapid utilization of iron by LD-R inside LD-R-PV to support its higher proliferation. To determine the key players that could drive the rapid mobilization of the excess labile iron pool into PV of LD-R infected MФs as compared to LD-S infected MФs, we looked into several well-reported iron regulatory, metabolizing, and transporter proteins by performing a comparative RNAseq analysis between LD-R and LD-S infected MФs 4hrs and 24hrs pi as compared to uninfected control. Transcriptomics analysis showed a significant change in a large number of transcripts in response to LD-R and LD-S infected MФs at 4hrs and 24hrs pi as compared to uninfected control as represented by the volcano plot (**Supplementary Figure 2B**). However, no significant change in the transcript expression of iron metabolizing and transporters like *trf* (CD71, Transferrin receptor 1), *slc40a1* (Ferroportin), *slc11a1* (NRAMP1), *slc11a2* (DMT1) were observed between LD-S and LD-R infected MФs as represented by a snapshot of all iron-related genes depicted in heat map (**Supplementary Figure 2C**). It should be noted that previous transcriptomic analysis to find altered transcripts of iron-regulated machinery in LD-infected MФs by others has also revealed no significant change (43). And previous reports have also pointed out that LD infection can regulate the expression of host iron-regulatory mechanism by post-transcriptional/post-translational modifications and this has been already reported for iron-regulating genes like NRAMP1, Transferrin receptor 1, Ferritin, and Ferroportin (42–45). Hence, the expression of a few targets that might help in iron scavenging inside LD-PV was checked at the protein level. Co-localization and Western blot analysis of CD71 (Transferrin receptor 1) with early endosome harboring LD-S and LD-R within infected MФs do not reflect any significant change in the level of CD71 between LD-S and LD-R infection, with early PV of both LD-S or LD-R showing significant expression of CD71 (**Supplementary Figure 3A.i. and A.ii**.) suggesting a basal level of iron needed for survival is provided by CD71 in both LD-S and LD-R. This observation is in concurrence with a previous report which also observed the same in the case of LD-infected MФs (43). Next, the expression pattern of Ferroportin was determined which is the sole known iron exporter that is expressed in macrophage membranes. Strikingly, a significant difference in localization and expression profile of the Ferroportin receptor in LD-R as compared to LD-S infected MФs was observed at 4hrs pi (**Figure 4C.i. and C.ii**.). A significant amount of Ferroportin was found to be colocalized with Rab5a containing early PV harboring LD-R as opposed to LD-S at 4hrs pi where Ferroportin is mostly restricted to MФs surface (**Figure 4C.i**.). Moreover, both LD-S and LD-R infection resulted in a significant increase in total Ferroportin expression at 4hrs pi as compared to uninfected control, although this increase appears to be slightly more in LD-R as compared to LD-S (**Figure 4C.ii**.). To prove that this selective re-localization of Ferroportin might help in iron mobilization inside LD-R-PV from host cytoplasm, MФs were pre-treated with Hepcidin, which degrades Ferroportin when exposed in MФs surface and then infected with LD-S or LD-R. Super-resolution microscopy of Hepcidin pre-treated and post-treated conditions for LD-S infected MФs showed no significant accumulation of Ferroportin around LD-S for either of the conditions (**Figure 4D.i. and Supplementary Figure 3C**). However, for LD-R infection, while no Ferroportin was observed around LD-R in Hepcidin pre-treatment, post-treatment of Hepcidin did result in significant accumulation of Ferroportin around LD-R, clearly suggesting that LD-R infection reshuffles Ferroportin from host plasma membrane to its PV rendering Hepcidin-mediated degradation of Ferroportin ineffective in already LD-R infected MФs (**Figure 4D.i., lowermost right panel)**. Simultaneous live cell imaging with Calcein-AM and lysotracker Red revealed the disappearance of ferroportin in Hepcidin pre-treated MФs fails to pump iron inside LD-R-PV as observed by higher calcein fluorescence similar to LD-S infection (**Figure 4D.ii., upper panel**). On the contrary, hepcidin post- treatment results in a rapid reshuffling of ferroportin around the PV membrane which also corroborates with rapid iron trafficking inside LD-R-PV as observed by reduced Calcein signal inside LD-R-PV, while hepcidin pre- and post-treatment have no bearing on LD-S-PV (**Figure 4D.ii., lower panel**). It was also interesting to note that NRAMP1 which has been previously implicated in the process of iron export from *L. major* infected PV and which also reflected no significant change in transcript level between LD-S and LD-R infection (**Supplementary Figure 2C**), was found to be present with slightly reduced expression around the early PV of LD-R as compared to early PV harboring LD-S, although no significant change in total NRAMP1 level was observed between LD-S and LD-R infected MФs in Western blot, suggesting that NRAMP1 might be more selectively excluded from LD-R-PV (**Figure 4E.i. and E.ii**.). These results altogether suggested that both the iron scavenging and iron withholding capacity of LD-R-PV is much more in LD-R-PV than LD-S-PV and is mainly regulated by post-transcriptional reprogramming of host MФs.

**Figure 4:**
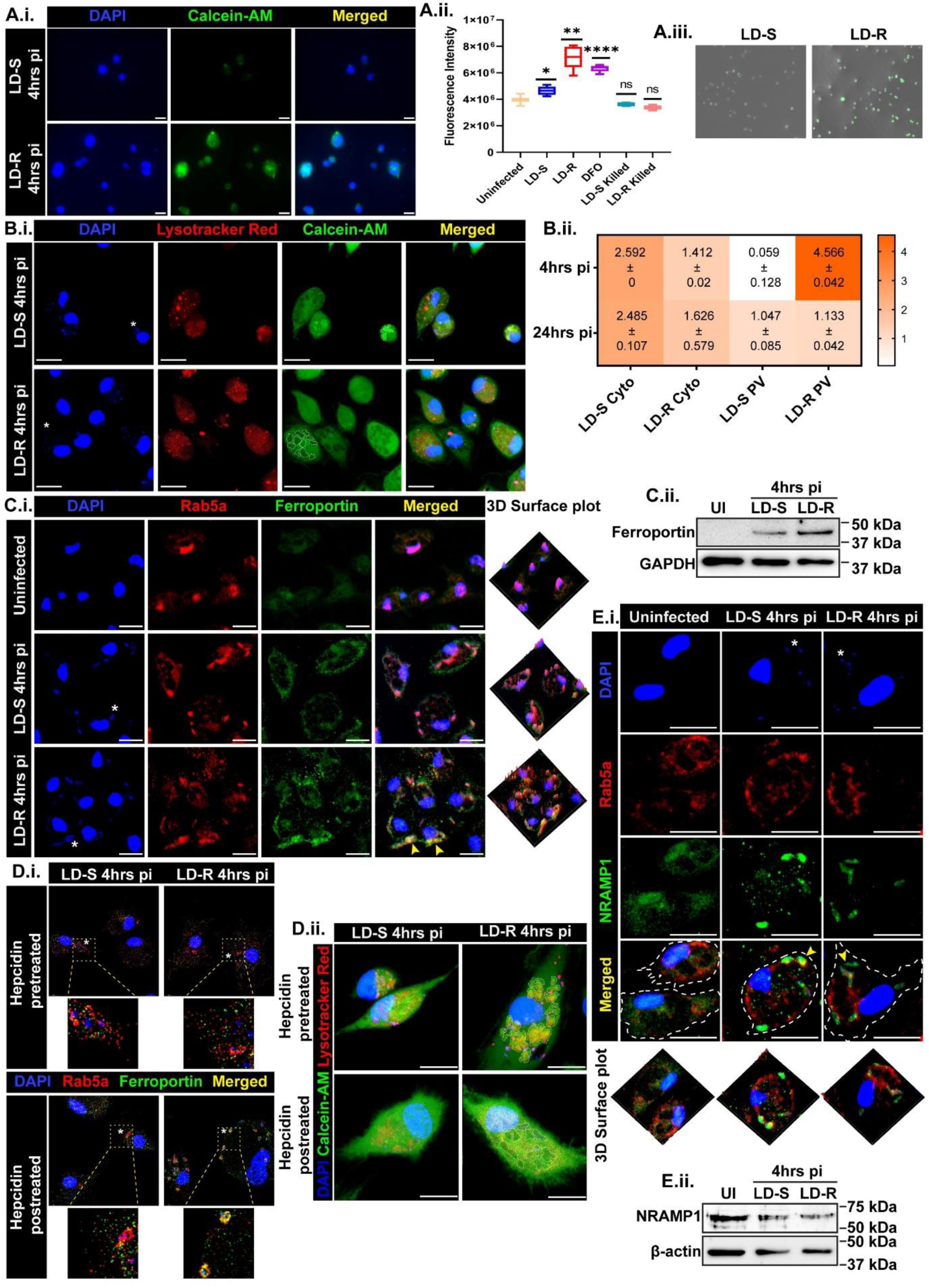
Reshuffling of Ferroportin and Nramp1 around LD-R-PV acts as a strategy to rapidly scavenge iron. **(A.i.)** Live-cell microscopy images of LD-S (upper panel) and LD-R infected MФs (lower panel) at 4hrs pi by staining with Calcein-AM which fluoresces (green) in low iron and gets quenched in the presence of iron. DAPI (blue) stains the nucleus. **(A.ii.)** Box plot showing fluorescence intensity of Calcein as in uninfected, LD-S, LD-R 4hrs pi, 100µM DFO treated (iron chelator), LD-S killed and LD-R killed 4hrs pi. **(A.iii.)** One representative field of LD-S (left) and LD-R (right) 4hrs pi in 10X showing Calcein fluorescence merged with DIC. **(B.i.)** Live-cell confocal microscopy of MФs infected with LD-S (upper panel) and LD-R (lower panel) at 4hrs pi stained with Calcein-AM and Lysotracker Red (red, which labels the PV). PV is marked in dotted areas in the 3^rd^ panel). **(B.ii.)** Heatmap showing quantification of iron in cytoplasmic fraction and PV fraction of LD-S and LD-R infected MФs at 4hrs (upper panel) and 24hrs pi (lower panel) following Ferrozine-based colorimetric assay using FeCl_3_ as standard. Data are represented as Mean ± SEM **(C.i.)** Confocal images showing Ferroportin localization in uninfected, LD-S, and LD-R infected MФs at 4hrs pi by labeling with anti-Ferroportin antibody (green) and anti-Rab5a antibody (Red). Yellow arrows in the merged panel of LD-R 4hrs pi show colocalization of Ferroportin with Rab5a and DAPI. The rightmost panel shows the 3D surface plots of representative sets. **(C.ii.)** Western blot of whole cell lysate of 3 experiment sets as mentioned in D.i. showing Ferroportin expression highest in LD-R at 4hrs pi with GAPDH as the loading control. **(D.i.)** Super-resolution confocal images of MФs either pretreated with hepcidin (1µg/ml) followed by LD infection for 4hrs or LD infection followed by hepcidin posttreatment and subsequently labeled with anti-Ferroportin in green and anti-Rab5a in red (which demarcates early PV) antibody and DAPI (marking the nucleus) before Confocal imaging. The left panel shows LD-S 4hrs pi and the right panel shows LD-R 4hrs pi with enlarged views of the selected region at the bottom of each set. Yellow fluorescence in the merged image denotes perfect colocalization of Rab5a, Ferroportin, and DAPI which is typically observed in MФs infected with LD-R for 4hrs followed by hepcidin posttreatment (lowermost right panel). **(D.ii.)** Live cell confocal images showing the status of the iron level in similar experimental sets D.i. staining with Calcein-AM and Lysotracker Red. The dotted region shows the PV portion with distinct changes. **(E.i.)** Confocal images showing expression of NRAMP1 (green) around PV, stained by anti-Rab5a antibody (Red), of LD-S and LD-R infected MФs at 2hrs pi. The lowermost panel shows the 3D surface plots of representative experimental sets. **(E.ii.)** Western blot of whole cell lysate of uninfected, LD-S 2hrs pi, and LD-R 2hrs pi showing NRAMP1 expression lowest in LD-R at 2hrs pi keeping β-actin as the loading control. One representative small nucleus of LD has been marked in (**\***) to show the infected MФs. The scale bar indicates 20µm. Each experiment was performed in 3 biological replicates and graphical data were represented as Mean with SEM. P > 0.05 is marked as ‘ns’ (non-significant), P ≤ 0.05 is marked as *, P ≤ 0.01 is marked as **, and P ≤ 0.0001 is marked as ****.

### 5. Iron depletion in the cytoplasm of LD-R infected macrophages quenches excess ROS through non-canonical activation of NRF-2 at early hours

Excess trafficking of iron inside LD-R-PV should result in the depletion of iron in the cytoplasm as compared to LD-S infection. Previous reports have suggested that low iron activates p62/SQSTM1, which in turn can degrade the NRF2 inhibitor Keap1 (46), allowing nuclear translocation of NRF2 and its subsequent binding with anti-oxidant response element (ARE)-promoter to activate the anti-oxidative pathway. Thus, iron deprivation could be the primary reason for bringing down elevated levels of ROS at a later time point in LD-R infected MФs (**Figure 2A.i. and A.ii**.). A significantly higher expression of p62 in the cytoplasm of LD-R infected MФs similar to DFO treated MФs as compared to LD-S infection at 4hrs pi further enforced this observation (**Supplementary Figure 4A.i. and A.ii**.). Higher p62 expression at 4hrs pi also corroborated with increased NRF2 expression in the whole cell lysate and nuclear fraction of LD-R infected MФs at 4hrs pi as compared to LD-S infected or uninfected MФs at the same time point (**Supplementary Figure 4B.i.a. and B.i.b**.). Furthermore, confocal microscopy with RGB-profile plot coupled with super-resolution 3-D rendering Z-stack images at 4hrs pi, also revealed a significant translocation of NRF2 in the nucleus in response to LD-R infection, while NRF2 is largely retained in the cytoplasm of LD-S infected MФs (**Supplementary Figure 4B.ii. and B.iii**.). These observations indicate LD-R infection results in reduced iron levels in the host cytoplasm around 4hrs pi leading to p62-mediated activation of NRF-2 which can lower ROS levels at a later time point.

### 6. Iron insufficiency in later time points is compensated by inflated erythrophagocytosis driven by SIRPα degradation

Previously Ferrozine-based colorimetric quantification of iron in cytoplasmic and PV fraction of LD-R infected MФs (**Figure 4B.ii**.) showed a slight but significant recovery in cytoplasmic iron level 24hrs pi. It was suspected that a cytoplasmic drop in the iron level is prompting LD-R infected MФs to uptake more iron from the medium to continue its rapid proliferation. Conventionally, the recycling of dead phagocytosed RBCs into iron is the major contributor of iron sources in MФs (47). A comparative analysis of erythrophagocytosis between LD-R and LD-S infected live MФs at 4hrs and 24hrs pi, showed, that although there is no significant difference in RBC uptake between LD-R and LD-S infected MФs at 4hrs pi (**Figure 5A.i. and A.ii**.), however at 24hrs pi, LD-R infected MФs specifically showed a huge accumulation of ingested RBC significantly co-localizing with GFP-positive LD (**Figure 5A.iii. and A.iv**.). Fluorometric plate reader-based quantification of erythrophagocytes at 4hrs and 24hrs pi also showed a significantly higher fluorescence intensity for CytoRed RBC in LD-R infected MФs specifically at 24hrs pi, while MФs incubated with LD-S or LD-R behaves similarly as uninfected control (**Figure 5A.ii. and A.iv**.). Signal regulatory protein α (SIRPα), a specific receptor on the MФ surface that interacts with CD47-enriched live RBCs to prevent erythrophagocytosis, was found to be significantly degraded on the surface of LD-R infected MФs at 24hrs pi, while a similar level of SIRPα observed in both LD-S and LD-R infected MФs 4hrs pi (**Figure 5B.i**.). Western blot of whole cell lysate confirmed the previous observation of SIRPα getting degraded in LD-R infected MФs at 24hrs pi as compared to 4hrs pi (**Figure 5B.ii**.), while for LD-S infection there was an increased expression of SIRPa at 24hrs. Furthermore, Flow-cytometry based quantification of the SIRPα-positive population also revealed a significant drop in the SIRPα-positive population in LD-R infected MФs at 24hrs pi as compared to 4hrs pi, while for LD-S infected MФs there is an increase in SIRPα-positive population (**Figure 5B.iii**.). Contrarily, CD11b, a raft-associated membrane marker that has been reported to show diminished expression upon LD infection (48–50), was found to be significantly low for both LD-S and LD-R infected MФs (**Figure 5B.iii**.). Notably, erythrophagocytosis has been previously reported to be a late consequence of LD infection by post-translational downregulation of SIRPα on the surface of infected MФs (11).

**Figure 5:**
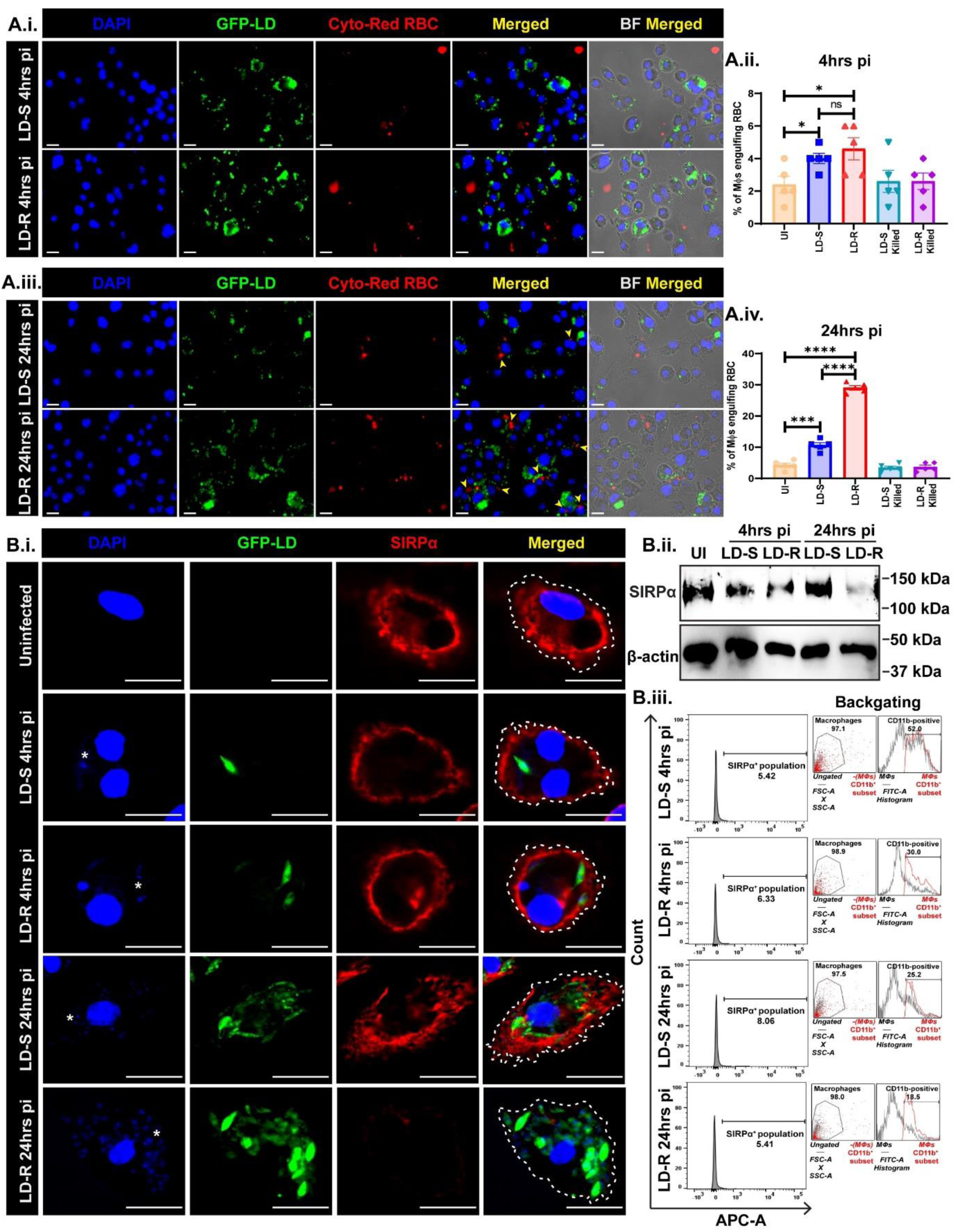
Augmented erythrophagocytosis in LD-R infected MФs at 24hrs pi in concomitance with SIRPα degradation. **(A.i.)** Confocal images show no significant erythrophagocytosed RBCs in both LD-S and LD-R infected MФs at 4hrs pi. DAPI stains nucleus, LD were labeled with GFP (green), and RBCs were labeled with Cyto-Red dye after infection to check engulfed RBCs in LD-infected MФs. **(A.ii.)** Bar graph showing % of MФs engulfing RBCs calculated from different fields of independent experimental triplicates at 4hrs pi. Killed-parasite infected MФs were kept as control. **(A.iii.)** Confocal images showing significant engulfed RBCs in LD-R infected MФs at 24hrs pi. The yellow arrow indicates ingested RBCs in infected MФs. **(A.iv.)** Bar graph showing % of MФs engulfing RBCs calculated from different fields of experimental triplicates sets at 24hrs pi. **(B.i.)** Confocal images showing SIRPα expression (red) at the macrophage surface outline demarcated by a dotted line in infected MФs with LD expressing GFP (green). One representative small nucleus of LD has been marked in (**\***) to show the infected MФs. **(B.ii.)** Western blot of whole cell lysate showing SIRPα expression in uninfected, LD-S, LD-R infected MФs at 4hrs pi and 24hrs pi. β-actin is used as a housekeeping control **(B.iii.)** Flow-cytometric data showing histogram plot representing SIRPα expression in CD11b-positive MФ population analyzed in FlowJo v10. The right-most panel shows the back-gating of individual experimental sets. The scale bar indicates 20µm. Each experiment was performed in 3 biological replicates and graphical data was represented as Mean with SEM. P > 0.05 is marked as ‘ns’ (non-significant), P ≤ 0.05 is marked as *, P ≤ 0.01 is marked as **, P ≤ 0.001 is marked as ***, and P ≤ 0.0001 is marked as ****.

### 7. Furin and ADAM10, acting consecutively result in the extracellular cleavage of SIRPα in LD-R infected macrophages

Our previous observation revealed that low iron promotes augmented erythrophagocytosis in concomitance with SIRPα degradation in LD-R infected macrophages at 24hrs pi which is promptly possible if the extracellular domain that senses CD47-enriched live RBC is degraded. There is increasing evidence that SIRPα can be shed from the macrophage surface by the action of several proteases like matrix metalloproteinase 9 (MMP-9), a disintegrin, and metalloprotease 10 and 17 (ADAM10 and ADAM17). A recent report suggests that GI254023X, an ADAM10 inhibitor can inhibit extracellular SIRPα cleavage significantly from LD-infected MФs surface (11). The expression level of the activated form of ADAM10 was monitored between LD-R and LD-S infected MФs. While, an enriched precursor form of ADAM10 (~90kDa) was observed in the case of LD-S infected MФs, an increased mature active form of ADAM10 (~68kDa) was enriched in LD-R infected MФs 24hrs pi (**Figure 6A)**. Prior reports in other disease models suggest that ADAM10 gets activated by Furin (51), a proprotein serine convertase, which gets activated in iron-deprived conditions through post posttranslational maturation (52). Furin undergoes several posttranslational maturation steps starting from the Endoplasmic reticulum (ER) to the cis-Golgi network to the trans-Golgi network to achieve its active form which can shuffle between trans-Golgi and plasma membrane or can get shed as an active enzyme (53). Surprisingly, a higher Furin expression was observed in the whole cell lysate of LD-S infected MФs at 24hrs pi as compared to LD-R infected MФs (**Figure 6B.i**.). Interestingly, however, immunofluorescence-based assays pointed out that the majority of the Furin in LD-S infected MФs are localized around the endosomal compartment having a perinuclear localization similar to uninfected MФs which is possibly the inactive form (**Figure 6B.ii**.). In LD-R infected MФs, Furin is found to have a more diffused localization throughout the cytoplasm inclining towards the cell membrane, explaining the western blot data which probably suggests that activation of Furin probably leads to its release from the LD-R infected MФs. Western blot analysis of supernatant derived from uninfected, LD-S infected, and LD-R infected MФs showed increased Furin level in LD-R infected MФs 24hrs pi similar to DFO, (iron chelator) treated control (**Figure 6B.iii**.), which further confirmed that Furin sheds from LD-R infected MФs at 24hrs pi probably acting over membrane-prodomain of ADAM10 leading to its activation (**Figure 6B.iii**.). Furthermore, in/out co-localization assay by differential permeabilization, clearly showed that for LD-S infected MФs, a significant amount of Furin has perinuclear localization along with slight membrane localization, for LD-R infected MФs it is entirely localized in the membrane colocalizing with ADAM-10 (**Figure 6C**). Taken together these results suggest that low iron in LD-R infected MФs at 24hrs pi promotes Furin maturation resulting in shedding as an active enzyme that acts over ADAM10 prodomain cleavage and subsequent activation (m-ADAM10). m-ADAM10 cleaves SIRPα at the extracellular domain (**Figure 6D)** thus initiating erythrophagocytosis.

**Figure 6:**
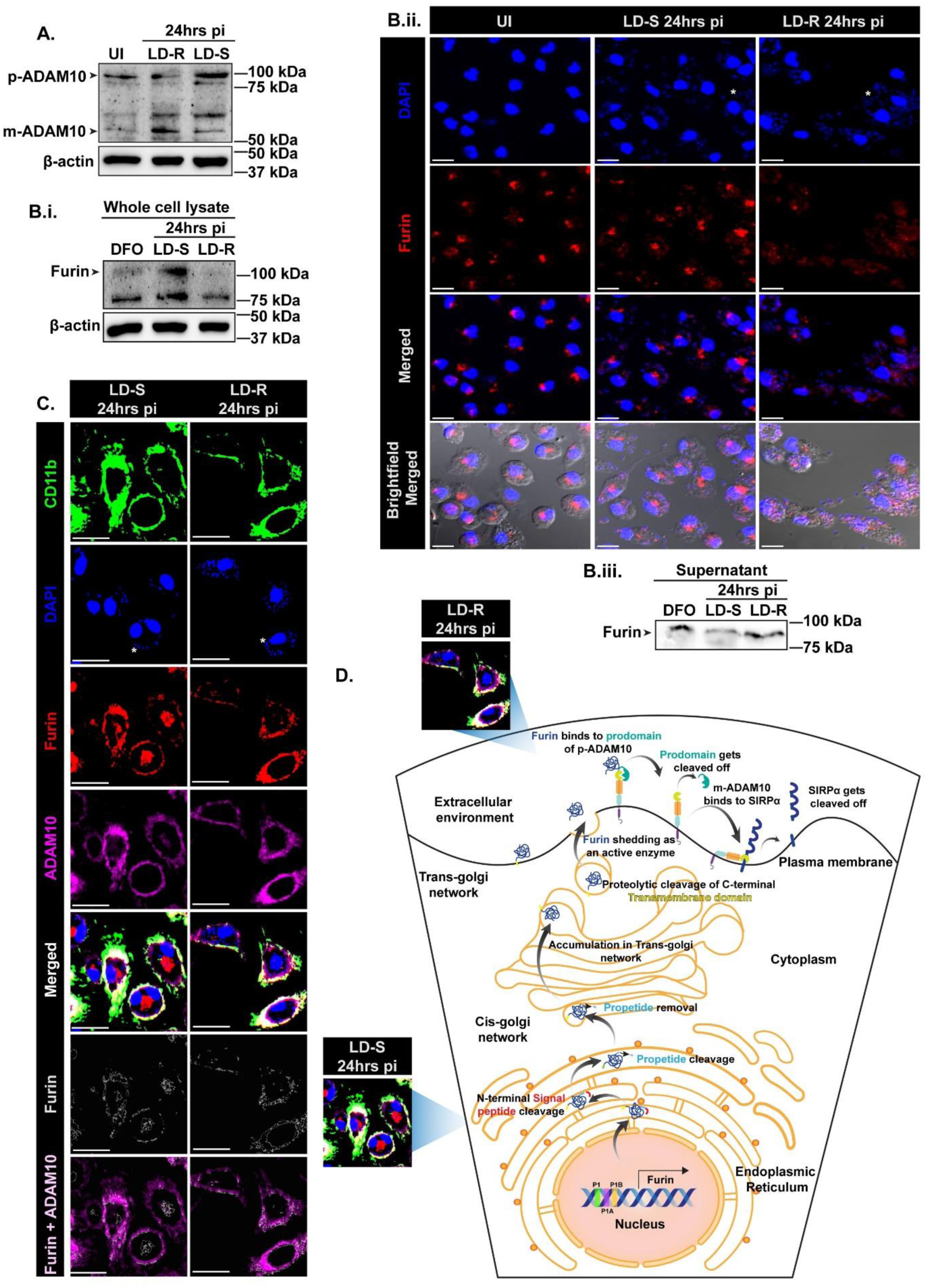
Low iron activates Furin which sheds to cleave the prodomain of ADAM10 rendering it activated. **(A.)** Western blot of whole cell lysate of uninfected, LD-R and LD-S infected MФs at 24hrs pi showing the expression of precursor-ADAM10 (p-ADAM10, ~90 kDa) and mature-ADAM10 (m-ADAM10, ~64 kDa) keeping β-actin acts as a housekeeping control. **(B.i.)** Western blot of whole cell lysate showing furin expression of DFO treated MФs, LD-S, and LD-R infected MФs at 24hrs pi. **(B.ii.)** 40X Confocal images showing the localization of furin (red) in uninfected, LD-S, and LD-R infected MФs. **(B.iii.)** Western blot of furin expression in the supernatant fraction of same experiments sets as B.i. **(C.)** Confocal images showing Furin (red, 3^rd^ panel and edges,6^th^ panel), ADAM10 (purple), and CD11b (green) colocalization in LD-S and LD-R infected MФs at 24hrs pi. One representative small nucleus of LD has been marked in (**\***) to show the infected MФs. The scale bar indicates 20µm. **(D.)** Scheme showing Furin maturation starting from Endoplasmic reticulum (ER) where it remains inactivated (observed in LD-S infected MФs at 24hrs pi) to getting shed as an active enzyme (observed in LD-R infected MФs at 24hrs pi) to cleave prodomain of p-ADAM10 rendering it activated (m-ADAM10), which in turn cleaves extracellular domain of SIRPα in LD-R infected MФs at 24hrs pi.

### 8. Infection with LD-R results in severe anemia in mice by inducing enhanced erythrophagocytosis

The major symptoms associated with VL are hepatosplenomegaly and anemia both of which are interlinked since an enlarged liver and spleen might contribute to the destruction of both healthy and senescent RBC which is a portrayal of extravascular hemolytic anemia. Infection with the LD-R parasites has been reported to cause increased organ parasite load (29) with a significantly enlarged spleen as compared to infection with their sensitive counterpart for both human and animal infection (2). The ability of LD-R to induce significantly higher erythrophagocytosis as compared to LD-S *in vitro* (**Figure 5A.iii. and A.iv**.) was further investigated in an experimental murine model of leishmaniasis as a possible reason for severe anemia. 4-6 weeks (N=3) old mice were infected with LD-S and LD-R in the tail vein or left uninfected and sacrificed at 24 weeks pi to evaluate comparative erythrophagocytosis as reported earlier (13). Although, hematoxylin-eosin (HE) stained splenic sections of uninfected, LD-S infected mice at 24 weeks pi revealed little or no significant erythrophagocytes, LD-R infected spleen showed significantly higher erythrophagocytes (**Figure 7A.i. and A.ii**.), with an increased percentage of phagocytosed RBCs being surrounded by LD-R amastigotes (**Figure 7A.iii**.). Hematology screening in an Erba H560 Fully Automated 5-part Hematology analyzer with emphasis on RBC-associated parameters performed for the investigation of anemia (**Figure 7B.i**.) revealed a drop in RBC and hemoglobin (HGB) count in LD-R-infected blood as compared to LD-S-infected blood. Importantly, a critically low hematocrit value (HCT) in LD-R-infected blood also confirmed severe anemia as compared to LD-S-infected blood. The other RBC indices (MCV, MCH, MCHC) do not show any significant difference between LD-S and LD-R mice, indicating that the mice in both these categories are physiologically similar; and free from any other chronic disease or nutritional deficiency disorder that could contribute to anemia. The similar RDW in both these categories also rule out the possibility of difference in the volume and size of the RBCs in both the groups thus emphasizing the fact that the drop of HGB, RBC count, and HCT in LD-R mice was due to the infection only. WBC-differential scatter plots via Low angle scatter (LS) vs medium angle scatter (MS) (**Figure 7B.ii**.) for uninfected, LD-S infected, and LD-R-infected blood showed enrichment of platelet aggregation or large RBC fragments in LD-R infected blood (marked as 2) as compared to LD-S infected or uninfected blood which might also indicate towards anemia severity. Other prominent regions marked as 8 i.e. monocytes, 9 i.e. neutrophils also revealed significant enrichment in LD-R as compared to LD-S or uninfected blood indicating inflammation resulting from persisting LD-R infection even at 24 weeks pi whereas fewer monocytes and neutrophils in LD-S point towards a resolving LD-S infection.

**Figure 7:**
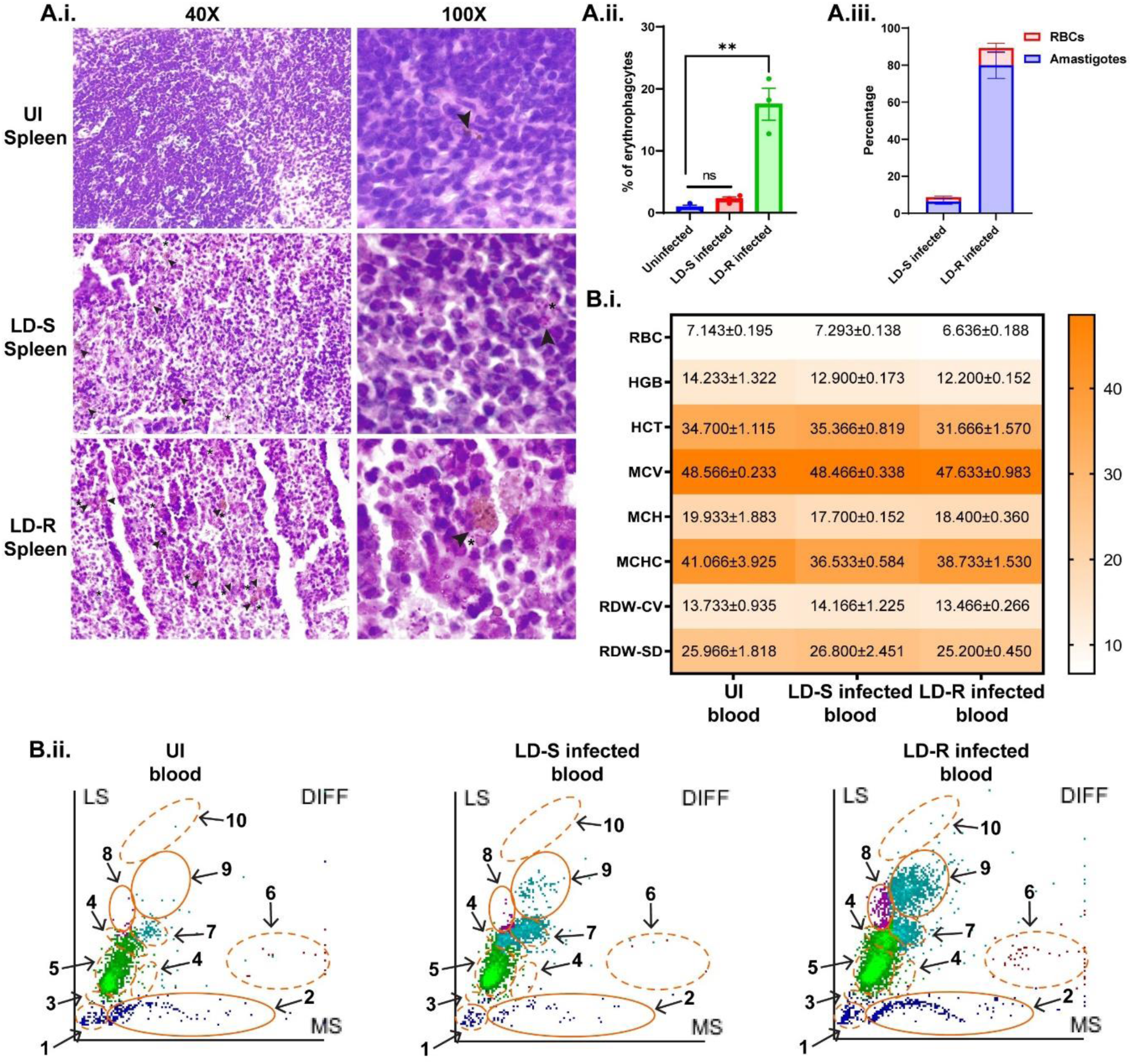
BALB/c mouse infected with LD-R for 24 weeks shows traits of severe anemia. **(A.i.)** Hematoxylin-eosin (HE)-stained splenic section of uninfected, LD-S infected, LD-R infected BALB/c mice (from upper panel to lower panel). The left panel shows 40X images showing a larger field of view and the right panel shows 100X images. The black arrow denotes ingested RBCs while a small nucleus of LD has been marked in (**\***) to show the infected MФs **(A.ii.)** Bar graph showing the total percentage of erythrophagocytes in uninfected, LD-S infected, and LD-R infected splenic sections. **(A.iii.)** Stacked bar plots showing % of amastigotes and RBC engulfed by erythrophagocytes in LD-S and LD-R infected spleen. **(B.i.)** Heatmap showing the differential count of RBC-associated parameters in uninfected, LD-S, and LD-R infected BALB/c mice. Data is represented as the Mean of experimental triplicates ± SEM. **(B.ii.)** WBC subpopulation Scatter plots of uninfected (left panel), LD-S infected (middle panel), and LD-R infected (right panel) BALB/c mice blood of one representative experimental set where the X-axis denotes the Medium angle scatter (MS) and Y-axis denotes the Low angle scatter (LS). Each subpopulation has been demarcated in (orange area) from 1-10. 1: Ghost, 2: Platelet aggregation or large RBC fragments, 3: Nucleated RBCs, 4: Abnormal lymphocytes, 5: Lymphocytes, 6: Eosinophils, 7: Left shift, 8: Monocytes, 9: Neutrophils, 10: Immature granulocytes. The dotted area signifies no significant change while the solid area denotes an area with significant changes. Each experiment was performed in 3 biological replicates and graphical data was represented as Mean with SEM. P > 0.05 is marked as ‘ns’ (non-significant) and P ≤ 0.01 is marked as **.

## Discussion

Widespread resistance to antimonial drugs has been reported in South Asia during the 1990s leading to higher rates of treatment failure ~60% (54). Hence antimonial drugs are inadvisable in India and Nepal. Nevertheless, LD isolated from clinical patients still show antimony resistance although antimony has been withdrawn from therapeutic interventions long back, suggesting a positive selection for antimony resistance in LD (1). In the Indian subcontinent rapid emergence of antimony resistance has been linked with the co-existence of the related element, arsenic in this region (55). Our findings revolve around understanding the mechanism by which these parasites might have evolved as a persistent strain in due course of evolution. In response to infection, mammalian hosts deploy an array of strategies to kill LD parasites and these parasites have evolved sophisticated yet diverse mechanisms to counteract the host defense strategy while proliferating inside host macrophages (MФs). In this regard, antimony-resistant LD (LD-R) has been reported to have a significantly higher parasite burden in the spleen and liver of infected humans and animals as compared to infection with their sensitive counterparts (LD-S) (2). Our observation pointed out that this high parasite burden linked with antimony resistance does not rely only on initial infectivity as commonly believed to date (2, 28) again indicating a selective survival advantage for antimony-resistant strains.

Since these parasites are predominantly antimony-resistant, to understand how these parasites evolved as a persister strain, the strategy for evading the anti-leishmanial mode of action of antimony should be decoded first. The mode of action of killing LD by pentavalent antimonials and initial host defense strategy includes the production of Reactive oxygen species (ROS) which causes DNA damage (56, 57). Logical interpretation leading to enhanced survival of antimony-resistant LD lines inside the host is their capability to prevent host-derived oxidative outbursts. Contrarily, LD-R instead of suppressing/ delaying ROS production, was found to elicit ROS production for its survival benefits. LD-R as opposed to LD-S were found to undergo a metabolic shift in their central carbon metabolism from Glycolysis to PPP resulting in enriched reducing intermediates allowing them to withstand higher oxidative stress (**Figure 2)**. Moreover, a previous report has also suggested that the presence of antimony-derived oxidative stress may result in the switching of central carbon metabolism towards PPP among intracellular LD (58). The fact that this induction of initial high ROS is observed only for LD isolates showing primary resistance to antimony but not for LD isolates showing unresponsiveness towards other drugs like Amp-B, probably indicates this might be a common occurrence among field isolates exhibiting antimony resistance. This posed an inherent question how is this initial oxidative outburst advantageous to LD-R in providing survival benefits? Of note, previous studies on *Trypanosoma cruzi* have also shown elevated ROS as a key signal for its proliferation inside host MФs (59, 60),. Our findings suggest that initial high ROS induces the overexpression of host Heme-oxygenase 1 (**Figure 3)** which degrades heme into the crucial bioavailable Fe^2+^ (61) which these LD parasites critically lack (62) and relies on host scavenging to ensure their survival. Iron acts as a critical element for several important biological processes including DNA replication, ATP production, mitochondrial respiration, etc. (63–65). Higher iron production and its subsequent mobilization within PV of LD-R thus satisfy their need for iron to support their rapid proliferation.

To date, most studies illustrated how LD parasites prevented iron efflux as a strategy to withhold iron (45) with little or no knowledge of understanding how iron/ heme/ hemoglobin is trafficked inside the PV, the residential niche for LD. However, the existence of iron, heme, and hemoglobin transporters has been well-reported in the *Leishmania* genome (62, 66, 67). Our observations clearly showed a higher abundance of Fe^2+^ in LD-R-PV (**Figure 4B)**, indicating a higher influx of host-derived iron specifically for LD-R amastigotes. However, equal distribution of CD71 (Transferrin receptor 1) around PV of both LD-R and LD-S infected MФs suggested the basal level of iron is provided by CD71 concerning both types of input infection (**Supplementary Figure 3A)**. Interestingly, our study pointed out overexpression and reshuffling of Ferroportin iron exporter around LD-R PV membrane during the formation of the phagosomal compartment as the critical component that allows LD-R to have more access to host iron (**Figure 4C)**. A previous report on *Salmonella typhimurium* has suggested an inward reorientation of Ferroportin is responsible for providing sufficient iron in *Salmonella*-containing vacuole rather than its conventional role in the exclusion of iron from infected host (68). Interestingly, Hepcidin, a peptide hormone produced in the liver, has been reported to degrade Ferroportin only when they are exposed to MФ surface (68). Reduced Fe^2+^ accumulation in LD-R-PV in the case of Hepcidin pre-treatment (**Figure 4D)** convincingly proved that LD-R has devised a specific strategy to reorient Ferroportin around intracellular LD-R, as compared to LD-S for which Ferroportin seems to have a minimal localization around the intracellular parasite (**Supplementary Figure 3B)**. Increased influx of iron inside LD-R-PV results in a drop of iron in the cytoplasm which in turn promotes non-canonical activation of NRF2 in the LD-R infected MФs leading to anti-oxidative response to bring down inflated ROS at a later time point around 8hrs pi (**Supplementary Figure 4, and Figure 2A)**. Another interesting observation was a surprised recovery of cytoplasmic Fe^2+^ level with a significant drop of iron in LD-R-PV at later time points around 24hrs pi (**Figure 4B)**. It can be safely assumed once most of the iron is utilized by LD-R, a continued supply of iron will be still required to maintain its ongoing proliferation. MФs are the key hub for recycling senescent RBCs to produce iron through a process termed erythrophagocytosis. Previous reports have shown significant erythrophagocytosis in LD-infected MФs and mice (27). These observations along with the clinical report of severe anemia in LD-R infected patients (69, 70), prompted us to investigate erythrophagoctosis as a possible cause of anemia in LD infection. Cross-talk between CD47 which is enriched and expressed in live RBCs and SIRPα which is expressed in MФ membrane is the critical determinant of erythrophagocytosis. When a live RBC carrying CD47 flag comes in contact with intact SIRPα, it elicits a ‘don’t eat me signal’ thus the macrophage discriminately phagocytoses dead or senescent RBCs leaving behind the live ones. Our results revealed significantly higher degradation of SIRPα on the surface of LD-R infected macrophages suggesting a breakdown of discrimination between live or dead RBCs and thus leading to augmented erythrophagocytosis (**Figure 5)**. Our results could also explain a possible connection between low cytoplasmic iron in LD-R infected MФs at 24hrs pi with subsequent loss of SIRPα. The low iron level has been previously reported to activate Furin (52), a proconvertase that reaches out to macrophage membrane and cleaves the prodomain of ADAM10, a disintegrin and metalloproteinase, to activate it (53). ADAM10 has been recently reported to promote extracellular proteolysis of SIRPα (71). Thus, connecting these dots, we were able to show that while for LD-S infected macrophages Furin was mostly restricted to a perinuclear localization, for LD-R infected ones they are mostly present on the MФ surface, co-localizing with ADAM10 to activate it (**Figure 6)**, with SIRPα degradation and enhanced erythrophagocytosis being the outcome. Finally, our results comparing LD-S and LD-R infection in a murine model of VL-induced anemia validated a higher percentage of erythrophagocytosis in LD-R infected mice, with a higher frequency of ingested RBCs among LD-R amastigotes (**Figure 7)**. Blood parameters like RBC, HGB, and HCT also support this observation confirming a trait of severe anemia observed in LD-R-infected mice. Thus, this study deciphers downstream pathways starting from initial infection to showing aggressive pathogenesis leading to chronic hepatosplenomegaly and severe anemia in drug-resistant LD infection which was hitherto unexplored. Our findings throw light on refining therapeutic strategies that might help in treating clinical drug-resistant LD infection and should explain the rapid emergence of antimony resistance in *Leishmania donovani* and related organisms.

## Figures and legends

**Supplementary Figure 1:**
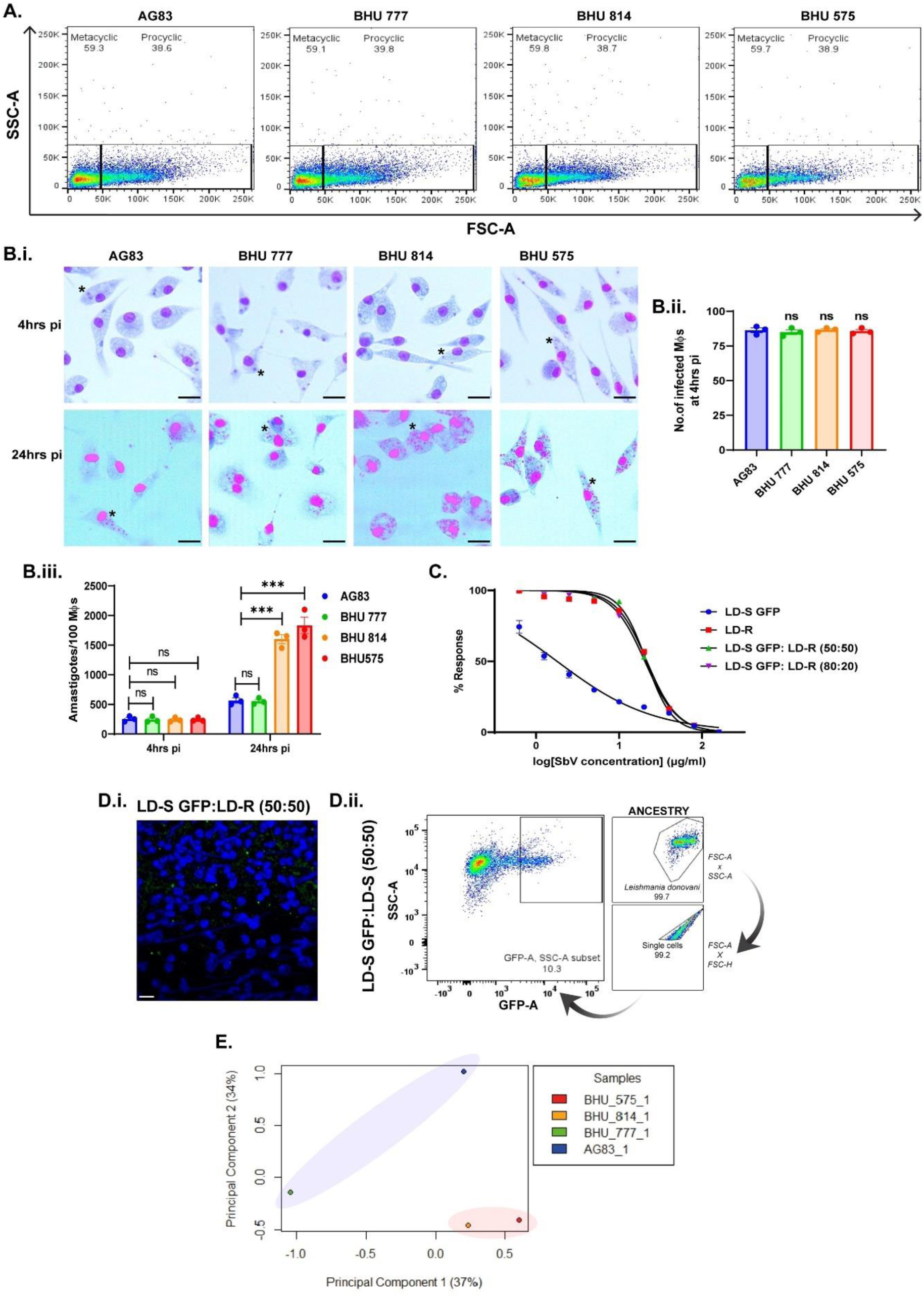
Antimony-resistant phenotype coincides with higher parasite burden if metacyclogenesis is kept as a constant factor. **(A.)** % of sorted metacyclic promastigote in AG83, BHU 777 (7^th^-day culture); BHU 814, BHU 575 (5thrd-day culture) by Beckman Coulter Cytoflex srt **(B.i.)** Giemsa-stained images of an equal number of AG83, BHU 777, BHU 814, and BHU 575 metacyclic promastigote-infected MФs at 4hrs and 24hrs pi. One representative small nucleus of LD has been marked in (*) to show the infected MФs. **(B.ii.)** No. of infected MФs/ 100 MФs at 4hrs pi were enumerated. Each data is the mean of three individual sets. **(B.iii.)** Amastigotes/100 MФs at 4hrs and 24hrs pi were calculated taking different fields and the data represents the SEM of three independent experiments. **(C.)** The dose-response curve of clonal LD lines derived from LD-S GFP, LD-R, LD-S GFP: LD-R (50:50), and LD-S GFP: LD-R (80:20) infected spleen to SbV as determined from intracellular amastigotes/100 MФs for calculating EC50. **(D.i.)** Confocal images of macerated spleen sample of LD-S GFP: LD-S showing the GFP amastigote load. The scale bar indicates 20µm. **(D.ii.)** % of the GFP-positive population was enumerated from FACS to denote the load of LD-S GFP. The right-most panel shows the ancestry of each analysis. **(E.)** PCA plot showing two separate clustering of LD-S (BHU 777 and AG83) demarcated in blue and LD-R (BHU 575 and BHU 814) demarcated in red. P > 0.05 is marked as ‘ns’ (non-significant) and P ≤ 0.001 is marked as ***.

**Supplementary Figure 2:**
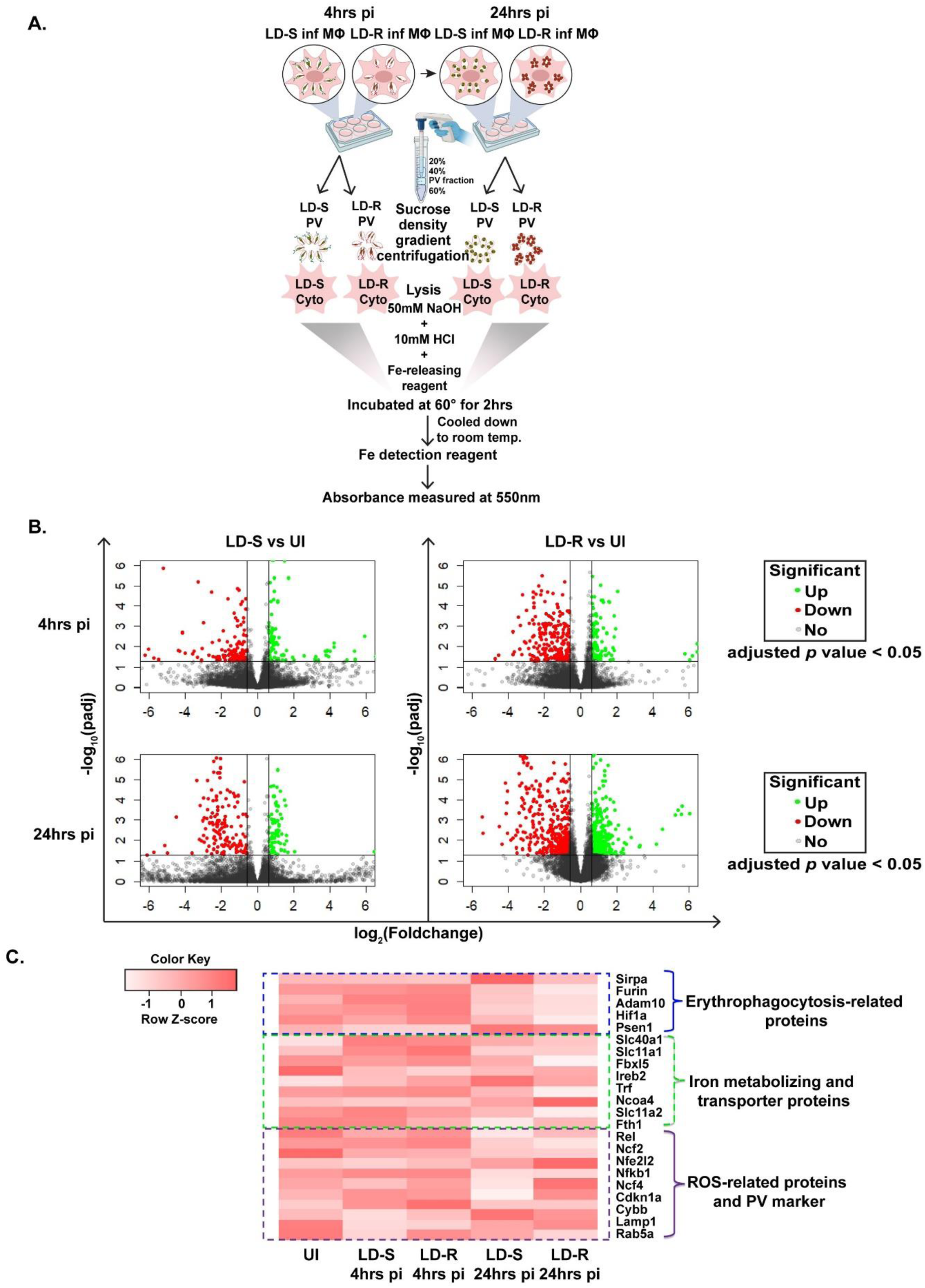
Schematic representation of Ferrozine-based iron quantification of PV and cytoplasmic fraction and transcriptomic landscape of LD-S and LD-R infected MФs at 4hrs and 24hrs pi as compared to uninfected MФs. **(A.)** Scheme showing PV extraction from LD-S and LD-R infected MФs at 4hrs and 24hrs pi ensued by quantification of iron following Ferrozine-based colorimetric assay. **(B.)** Volcano plots of differential gene expression of LD-S (left) and LD-R (right) infected MФs versus uninfected control at 4hrs pi (upper panel) and 24hrs pi (lower panel). Genes above the significance threshold (adjusted *p-value*<0.05) are marked in green having log_2_(fold change)>0.6 i.e. upregulated, and red having log_2_(fold change)<-0.6 i.e. downregulated, while the rest are marked in grey. **(C.)** Heatmap showing the differential expression pattern of erythophagocytosis-related protein (demarcated in blue), iron-metabolizing and transporter protein (green), and ROS-related proteins (purple) in uninfected, LD-S, and LD-R infected MФs at 4hrs and 24hrs pi (p>0.5).

**Supplementary Figure 3:**
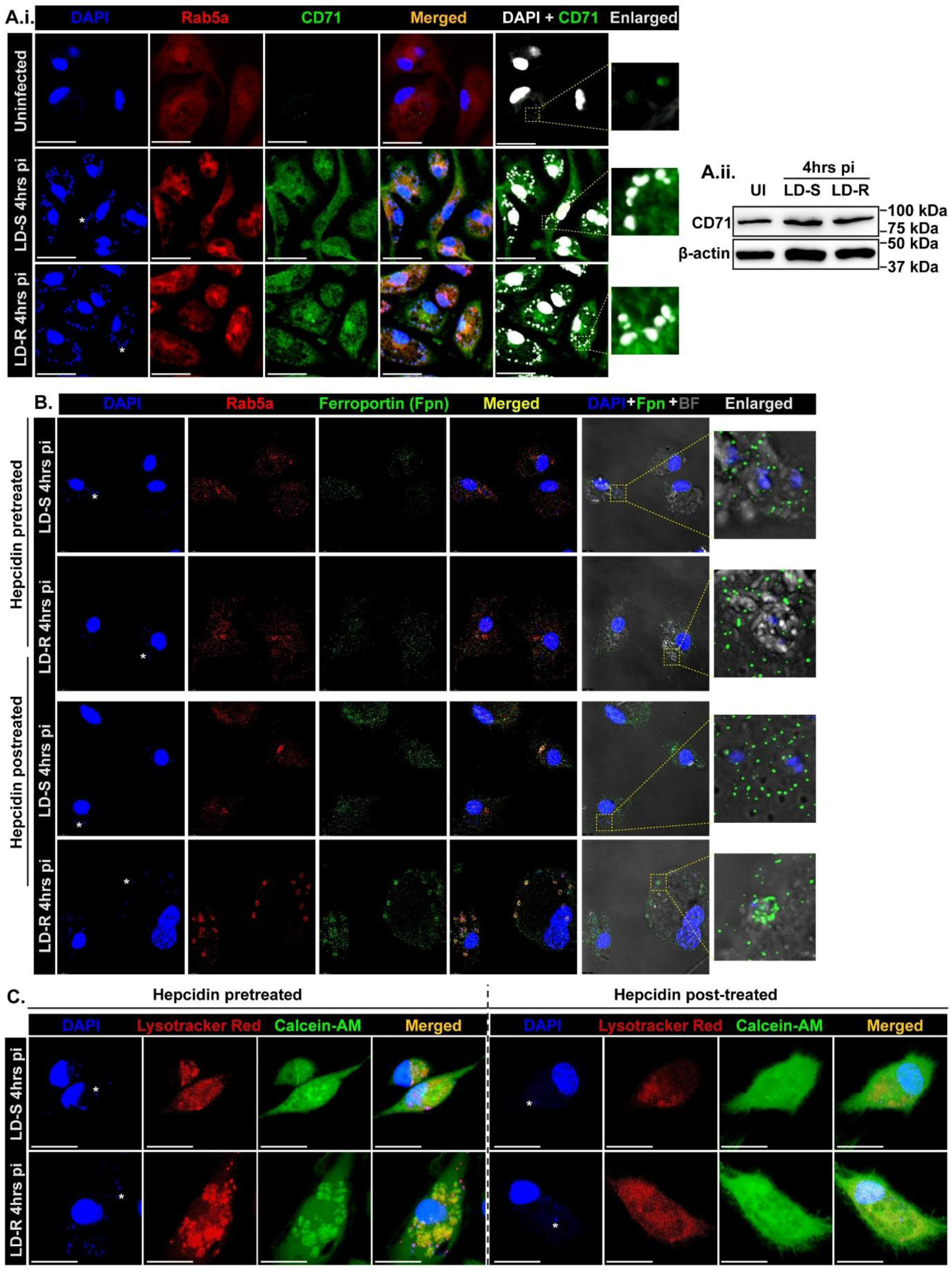
Localization of Ferroportin and CD71 around LD-PV. **(A.i.)** Confocal images showing the status of CD71 (Transferrin receptor 1) expression (green) in LD-S (upper panel) and LD-R infected MФs (lower panel) at 4hrs pi. Rab5a (red) demarcates early PV. **(A.ii.)** Western blot showing expression of CD71 in the whole cell lysate of uninfected, LD-S, and LD-R infected MФs at 4hrs pi. **(B.)** All panels of confocal images of hepcidin pretreated and hepcidin post-treated sets from Figure 4D.i. showing Ferroportin (green) localization. The right-most panel shows the enlarged view of a portion showing Ferroportin (green) localization with the LD nucleus (small blue dot) **(C.)** All panels of live cell confocal images showing iron status by staining with Calcein-AM (green) with lysotracker Red that demarcates PV in hepcidin pretreatment (left 4 panels) and hepcidin post-treatment (right 4 panels) experimental sets (separated by dotted line) elaborated from Figure 4D.ii. One representative small nucleus of LD has been marked in (**\***) to show the infected MФs. The scale bar indicates 20µm.

**Supplementary Figure 4:**
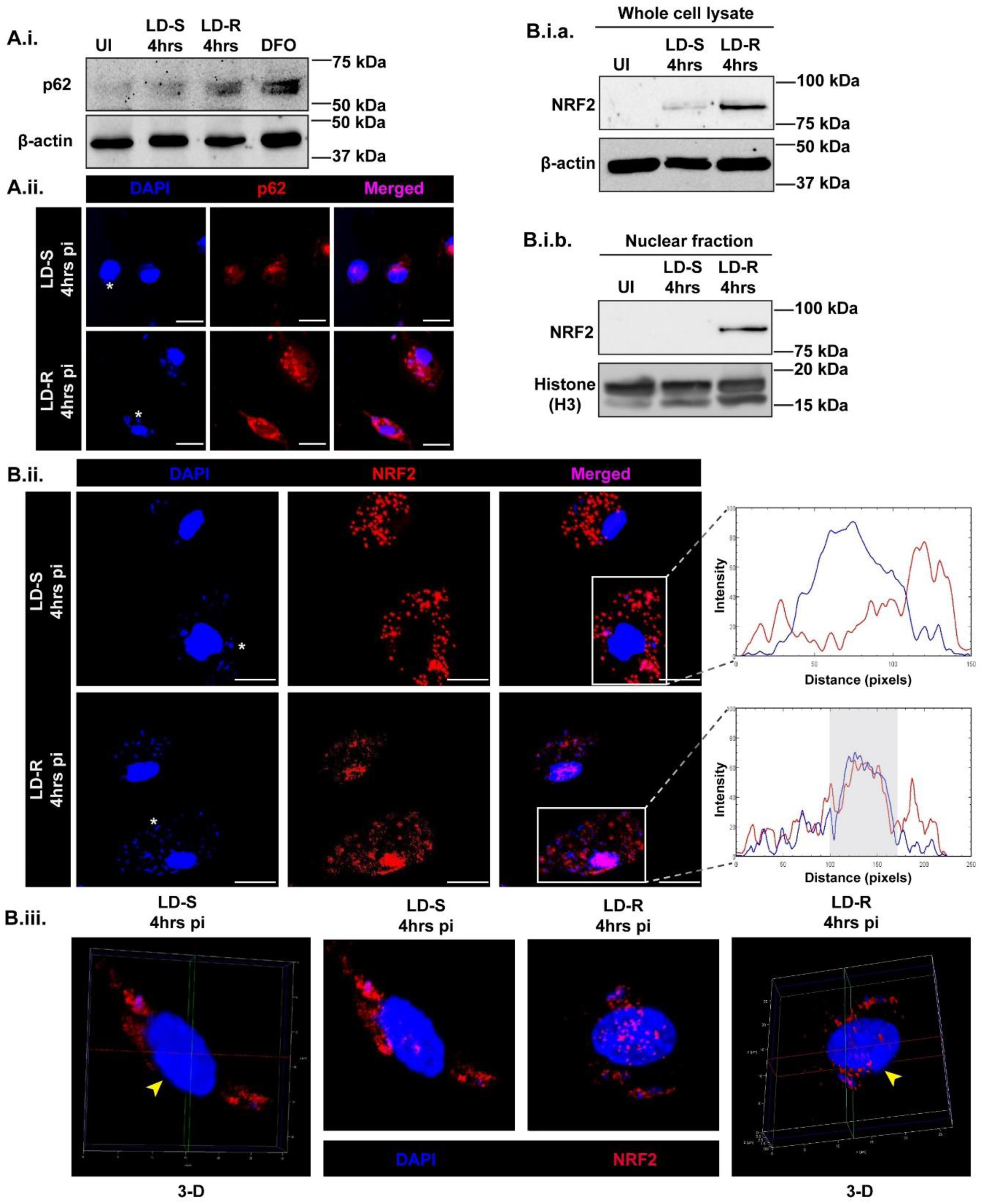
Low iron in the cytoplasm in LD-R infected MФs promotes p62/SQSTM1 mediated non-canonically activation of NRF2. **(A.i.)** Western blot of whole cell lysate showing the expression of p62 in uninfected, LD-S, LD-R infected MФs at 4hrs pi and DFO-treated MФs keeping β-actin as the loading control. **(A.ii.)** Confocal images showing the expression pattern of p62 (red) in LD-S and LD-R infected MФs at 4hrs pi. **(B.i.)** Western blot of whole cell lysate **(B.i.a.)** and nuclear fraction **(B.i.b.)** of uninfected, LD-S and LD-R infected MФs at 4hrs pi showing the expression level of NRF2 keeping β-actin and Histone (H3) as loading control respectively. **(B.ii.)** Confocal images showing the localization of NRF2 in LD-S and LD-S infected MФs at 4hrs pi. The left panel shows the RGB-profile plot of NRF2 (red) and DAPI (blue) where the X-axis denotes distance in pixels and the Y-axis denotes intensity. The grey area in the RGB-profile plot shows the colocalized region of NRF2 with nucleus **(B.iii.)** Super-resolution image showing NRF2 colocalization in the nucleus. The left-most and right-most panel shows 3D rendering of Z-stack images of LD-S and LD-R 4hrs pi respectively where the yellow arrow denotes a clipped quadrant in Leica Stellaris 5 image processing software to distinguish between superficially-present NRF2 and nuclear-localized NRF2. The middle two panels show the super-resolution Maximum intensity projection of Z-stack images of the respective sets showing the NRF2 (red) expression localization in the nucleus.

**Supplementary Table 1A:** Table showing the list of Glycolytic genes presented in the heatmap of Figure 2G.i.b with their respective enrichment scores.

| SI.No. | SYMBOL | TITLE | RANK IN GENE LIST | RANK METRIC SCORE | RUNNING ES | CORE ENRICHMENT |
| --- | --- | --- | --- | --- | --- | --- |
| 1 | LDBPK_361320 | Fructose-bisphosphate aldolase (EC 4.1.2.13) | 211 | 0.167 | 0.0559 | No |
| 2 | LDBPK_120490 | Glucose-6-phosphate isomerase (EC 5.3.1.9) | 232 | 0.16 | 0.1366 | No |
| 3 | LDBPK_141240 | Enolase | 1856 | 0.041 | 0.0089 | No |
| 4 | LDBPK_366960 | "2,3-bisphosphoglycerate-independent phosphoglycerate mutase" | 2433 | 0.027 | 0.0174 | No |
| 5 | LDBPK_362450 | Glucokinase | 3922 | -0.002 | -0.0928 | No |
| 6 | LDBPK_210700 | "Phosphoglucomutase, putative" | 5270 | -0.027 | -0.1846 | No |
| 7 | LDBPK_240870 | Triosephosphate isomerase (EC 5.3.1.1) | 6115 | -0.049 | -0.211 | Yes |
| 8 | LDBPK_041170 | "Fructose-1,6-bisphosphatase, cytosolic, putative" | 6365 | -0.058 | -0.16 | Yes |
| 9 | LDBPK_362480 | "Glyceraldehyde 3-phosphate dehydrogenase, cytosolic" | 6398 | -0.059 | -0.0808 | Yes |
| 10 | LDBPK_292620 | ATP-dependent phosphofructokinase | 6630 | -0.069 | -0.0275 | Yes |
| 11 | LDBPK_055450 | Pyruvate kinase (EC 2.7.1.40) | 6817 | -0.078 | 0.0316 | Yes |
| 12 | LDBPK_200110 | "Phosphoglycerate kinase C, glycosomal" | 6875 | -0.082 | 0.1075 | Yes |

**Supplementary Table 1B:**
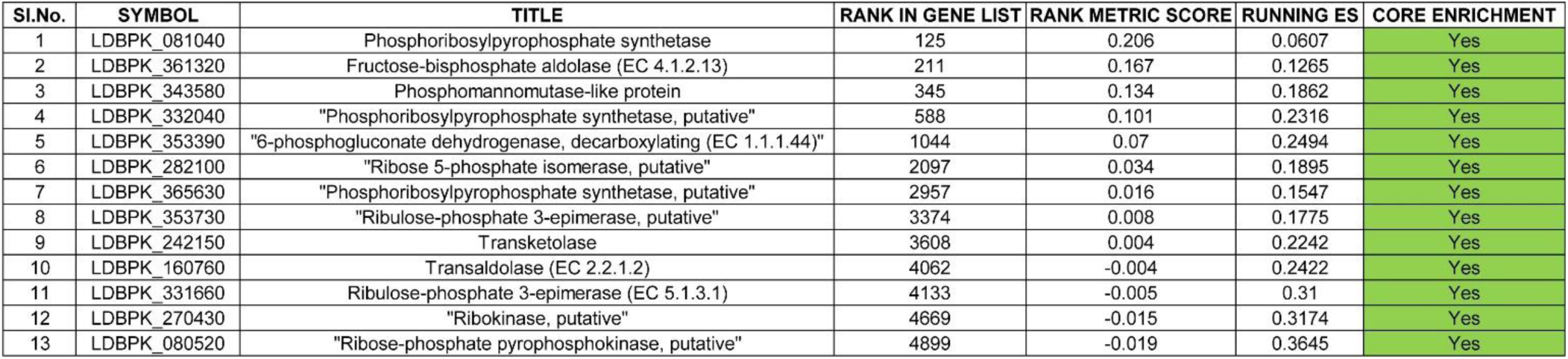
Table showing the list of Pentose phosphate pathway (PPP) genes presented in the heatmap of Figure 2G.ii.b with their respective enrichment scores.

## Acknowledgments

We acknowledge DST FIST, Govt. of India, for using the Confocal Microscope of the Biotechnology department, IIT Kharagpur for capturing most of the Confocal images. We acknowledge the Confocal Microscope purchased under the DST-FIST grant conferred on the School of Bioscience, IIT Kharagpur File No. SR/FST/LS-I/2019/595©. We acknowledge the Confocal Microscope of the Leica Stellaris 5 Dmi8 demonstration unit for capturing images of Figure 3. We would like to acknowledge the Central Research Facility of the School of Medical Science and Technology, IIT Kharagpur, India for providing access to instruments such as Flow Cytometer (BD Biosciences) and ERBA H560 Fully Automated 5-part Hematology Analyser. We would like to acknowledge Debolina Manna, Ph.D. scholar in IDI lab, SMST, IITKGP for generating transgenic LD-S GFP, LD-R GFP, and LD-R RFP lines.

## Author contributions

Conceptualization: B.M. and S.G.

Methodology Design: S.G., B.M., and Y.G. (erythrophagocytosis)

Formal Analysis: S.G.; K.V. & S.G. (Transcriptomics and Metabolomics); S.G. & S.D. (Histology)

Investigation: S.G.; K.V. & S.G. (Metabolomics-Figure 2H, Supplementary Figure 2B and 2C); S.G. & S.D. (Histology-Figure 7A); Y.G. and B.M. (Erythrophagocytosis-Figure 5A); B.M. (Iron Level-Figure 4A)

Writing-Original draft: S.G. and B.M.

Writing-Review and Editing: S.G., K.V., S.D., Y.G., B.M. Visualization: S.G., K.V., and B.M.

Supervision: B.M.

Project Administration and Reagents: B.M. and Y.G. Funding Acquisition: B.M., S.D., and Y.G.

All authors read and approved the final manuscript.

## References

1. Dumetz F, Cuypers B, Imamura H, Zander D, D’Haenens E, Maes I, et al. Molecular Preadaptation to Antimony Resistance in Leishmania donovani on the Indian Subcontinent. mSphere. 2018;3(2).

2. Vanaerschot M, De Doncker S, Rijal S, Maes L, Dujardin JC, Decuypere S. Antimonial resistance in Leishmania donovani is associated with increased in vivo parasite burden. PloS one. 2011;6(8):e23120.

3. Khan YA, Andrews NW, Mittra B. ROS regulate differentiation of visceralizing Leishmania species into the virulent amastigote form. Parasitol Open. 2018;4.

4. Rochael NC, Guimaraes-Costa AB, Nascimento MT, DeSouza-Vieira TS, Oliveira MP, Garcia e Souza LF, et al. Classical ROS-dependent and early/rapid ROS-independent release of Neutrophil Extracellular Traps triggered by Leishmania parasites. Scientific reports. 2015;5:18302.

5. Matte C, Arango Duque G, Descoteaux A. Leishmania donovani Metacyclic Promastigotes Impair Phagosome Properties in Inflammatory Monocytes. Infection and immunity. 2021;89(7):e0000921.

6. Bhardwaj S, Srivastava N, Sudan R, Saha B. Leishmania interferes with host cell signaling to devise a survival strategy. Journal of biomedicine & biotechnology. 2010;2010:109189.

7. Lodge R, Diallo TO, Descoteaux A. Leishmania donovani lipophosphoglycan blocks NADPH oxidase assembly at the phagosome membrane. Cellular microbiology. 2006;8(12):1922–31.

8. Huynh C, Andrews NW. Iron acquisition within host cells and the pathogenicity of Leishmania. Cellular microbiology. 2008;10(2):293–300.

9. Laranjeira-Silva MF, Hamza I, Perez-Victoria JM. Iron and Heme Metabolism at the Leishmania-Host Interface. Trends in parasitology. 2020;36(3):279–89.

10. Mahajan V, Marwaha RK. Immune mediated hemolysis in visceral leishmaniasis. J Trop Pediatr. 2007;53(4):284–6.

11. Hirai H, Hong J, Fujii W, Sanjoba C, Goto Y. Leishmania Infection-Induced Proteolytic Processing of SIRPalpha in Macrophages. Pathogens. 2023;12(4).

12. Varma N, Naseem S. Hematologic changes in visceral leishmaniasis/kala azar. Indian J Hematol Blood Transfus. 2010;26(3):78–82.

13. Morimoto A, Omachi S, Osada Y, Chambers JK, Uchida K, Sanjoba C, et al. Hemophagocytosis in Experimental Visceral Leishmaniasis by Leishmania donovani. PLoS neglected tropical diseases. 2016;10(3):e0004505.

14. Chakraborty P, Sturgill-Koszycki S, Russell DG. Isolation and characterization of pathogen-containing phagosomes. Methods Cell Biol. 1994;45:261–76.

15. Lu W, Wang L, Chen L, Hui S, Rabinowitz JD. Extraction and Quantitation of Nicotinamide Adenine Dinucleotide Redox Cofactors. Antioxidants & redox signaling. 2018;28(3):167–79.

16. t’Kindt R, Jankevics A, Scheltema RA, Zheng L, Watson DG, Dujardin JC, et al. Towards an unbiased metabolic profiling of protozoan parasites: optimisation of a Leishmania sampling protocol for HILIC-orbitrap analysis. Anal Bioanal Chem. 2010;398(5):2059–69.

17. Chambers MC, Maclean B, Burke R, Amodei D, Ruderman DL, Neumann S, et al. A cross-platform toolkit for mass spectrometry and proteomics. Nat Biotechnol. 2012;30(10):918–20.

18. Agrawal S, Kumar S, Sehgal R, George S, Gupta R, Poddar S, et al. El-MAVEN: A Fast, Robust, and User-Friendly Mass Spectrometry Data Processing Engine for Metabolomics. Methods in molecular biology (Clifton, NJ). 2019;1978:301–21.

19. Subramanian A, Tamayo P, Mootha VK, Mukherjee S, Ebert BL, Gillette MA, et al. Gene set enrichment analysis: a knowledge-based approach for interpreting genome-wide expression profiles. Proceedings of the National Academy of Sciences of the United States of America. 2005;102(43):15545–50.

20. Mootha VK, Lindgren CM, Eriksson KF, Subramanian A, Sihag S, Lehar J, et al. PGC-1alpha-responsive genes involved in oxidative phosphorylation are coordinately downregulated in human diabetes. Nat Genet. 2003;34(3):267–73.

21. Love MI, Huber W, Anders S. Moderated estimation of fold change and dispersion for RNA-seq data with DESeq2. Genome Biol. 2014;15(12):550.

22. Van Keuren-Jensen KR, Malenica I, Courtright AL, Ghaffari LT, Starr AP, Metpally RP, et al. microRNA changes in liver tissue associated with fibrosis progression in patients with hepatitis C. Liver Int. 2016;36(3):334–43.

23. Salvador-Martin S, Rubbini G, Vellosillo P, Zapata-Cobo P, Velasco M, Palomino LM, et al. Blood gene expression biomarkers of response to anti-TNF drugs in pediatric inflammatory bowel diseases before initiation of treatment. Biomed Pharmacother. 2024;173:116299.

24. Tenopoulou M, Kurz T, Doulias PT, Galaris D, Brunk UT. Does the calcein-AM method assay the total cellular ‘labile iron pool’ or only a fraction of it? The Biochemical journal. 2007;403(2):261–6.

25. Dighal A, Mukhopadhyay D, Sengupta R, Moulik S, Mukherjee S, Roy S, et al. Iron trafficking in patients with Indian Post kala-azar dermal leishmaniasis. PLoS neglected tropical diseases. 2020;14(2):e0007991.

26. Riemer J, Hoepken HH, Czerwinska H, Robinson SR, Dringen R. Colorimetric ferrozine-based assay for the quantitation of iron in cultured cells. Anal Biochem. 2004;331(2):370–5.

27. Morimoto A, Uchida K, Chambers JK, Sato K, Hong J, Sanjoba C, et al. Hemophagocytosis induced by Leishmania donovani infection is beneficial to parasite survival within macrophages. PLoS neglected tropical diseases. 2019;13(11):e0007816.

28. Ouakad M, Vanaerschot M, Rijal S, Sundar S, Speybroeck N, Kestens L, et al. Increased metacyclogenesis of antimony-resistant Leishmania donovani clinical lines. Parasitology. 2011;138(11):1392–9.

29. Vanaerschot M, Maes I, Ouakad M, Adaui V, Maes L, De Doncker S, et al. Linking in vitro and in vivo survival of clinical Leishmania donovani strains. PloS one. 2010;5(8):e12211.

30. Saha B, Pai K, Sundar S, Bhattacharyya M, Bodhale NP. The drug resistance mechanisms in Leishmania donovani are independent of immunosuppression. Cytokine. 2021;145:155300.

31. Roy M, Sarkar D, Chatterjee M. Quantitative monitoring of experimental and human leishmaniasis employing amastigote-specific genes. Parasitology. 2022;149(8):1085–93.

32. Moradin N, Descoteaux A. Leishmania promastigotes: building a safe niche within macrophages. Frontiers in cellular and infection microbiology. 2012;2:121.

33. Fernandez-Marcos PJ, Nobrega-Pereira S. NADPH: new oxygen for the ROS theory of aging. Oncotarget. 2016;7(32):50814–5.

34. Almugadam SH, Trentini A, Maritati M, Contini C, Rugna G, Bellini T, et al. Influence of 6-aminonicotinamide (6AN) on Leishmania promastigotes evaluated by metabolomics: Beyond the pentose phosphate pathway. Chem Biol Interact. 2018;294:167–77.

35. Mukhopadhyay R, Mukherjee S, Mukherjee B, Naskar K, Mondal D, Decuypere S, et al. Characterisation of antimony-resistant Leishmania donovani isolates: biochemical and biophysical studies and interaction with host cells. International journal for parasitology. 2011;41(13-14):1311–21.

36. Carneiro PP, Conceicao J, Macedo M, Magalhaes V, Carvalho EM, Bacellar O. The Role of Nitric Oxide and Reactive Oxygen Species in the Killing of Leishmania braziliensis by Monocytes from Patients with Cutaneous Leishmaniasis. PloS one. 2016;11(2):e0148084.

37. Paiva CN, Feijo DF, Dutra FF, Carneiro VC, Freitas GB, Alves LS, et al. Oxidative stress fuels Trypanosoma cruzi infection in mice. The Journal of clinical investigation. 2012;122(7):2531–42.

38. Andrews NW. Oxidative stress and intracellular infections: more iron to the fire. The Journal of clinical investigation. 2012;122(7):2352–4.

39. Silva R, Travassos LH, Paiva CN, Bozza MT. Heme oxygenase-1 in protozoan infections: A tale of resistance and disease tolerance. PLoS pathogens. 2020;16(7):e1008599.

40. Mukherjee B, Paul J, Mukherjee S, Mukhopadhyay R, Das S, Naskar K, et al. Antimony-Resistant Leishmania donovani Exploits miR-466i To Deactivate Host MyD88 for Regulating IL-10/IL-12 Levels during Early Hours of Infection. Journal of immunology. 2015;195(6):2731–42.

41. Riedelberger M, Penninger P, Tscherner M, Seifert M, Jenull S, Brunnhofer C, et al. Type I Interferon Response Dysregulates Host Iron Homeostasis and Enhances Candida glabrata Infection. Cell host & microbe. 2020;27(3):454–66 e8.

42. Banerjee S, Datta R. Leishmania infection triggers hepcidin-mediated proteasomal degradation of Nramp1 to increase phagolysosomal iron availability. Cellular microbiology. 2020;22(12):e13253.

43. Das NK, Biswas S, Solanki S, Mukhopadhyay CK. Leishmania donovani depletes labile iron pool to exploit iron uptake capacity of macrophage for its intracellular growth. Cellular microbiology. 2009;11(1):83–94.

44. Sen S, Bal SK, Yadav S, Mishra P, G VV, Rastogi R, et al. Intracellular pathogen Leishmania intervenes in iron loading into ferritin by cleaving chaperones in host macrophages as an iron acquisition strategy. The Journal of biological chemistry. 2022;298(12):102646.

45. Das NK, Sandhya S, G VV, Kumar R, Singh AK, Bal SK, et al. Leishmania donovani inhibits ferroportin translation by modulating FBXL5-IRP2 axis for its growth within host macrophages. Cellular microbiology. 2018;20(7):e12834.

46. Inoue H, Hanawa N, Katsumata SI, Katsumata-Tsuboi R, Takahashi N, Uehara M. Iron deficiency induces autophagy and activates Nrf2 signal through modulating p62/SQSTM. Biomed Res. 2017;38(6):343–50.

47. Klei TR, Meinderts SM, van den Berg TK, van Bruggen R. From the Cradle to the Grave: The Role of Macrophages in Erythropoiesis and Erythrophagocytosis. Frontiers in immunology. 2017;8:73.

48. De Almeida MC, Cardoso SA, Barral-Netto M. Leishmania (Leishmania) chagasi infection alters the expression of cell adhesion and costimulatory molecules on human monocyte and macrophage. International journal for parasitology. 2003;33(2):153–62.

49. Chakraborty D, Banerjee S, Sen A, Banerjee KK, Das P, Roy S. Leishmania donovani affects antigen presentation of macrophage by disrupting lipid rafts. Journal of immunology. 2005;175(5):3214–24.

50. Ghosh J, Das S, Guha R, Ghosh D, Naskar K, Das A, et al. Hyperlipidemia offers protection against Leishmania donovani infection: role of membrane cholesterol. J Lipid Res. 2012;53(12):2560–72.

51. Vassar R. ADAM10 prodomain mutations cause late-onset Alzheimer’s disease: not just the latest FAD. Neuron. 2013;80(2):250–3.

52. Silvestri L, Camaschella C. A potential pathogenetic role of iron in Alzheimer’s disease. J Cell Mol Med. 2008;12(5A):1548–50.

53. Braun E, Sauter D. Furin-mediated protein processing in infectious diseases and cancer. Clin Transl Immunology. 2019;8(8):e1073.

54. Olliaro PL, Guerin PJ, Gerstl S, Haaskjold AA, Rottingen JA, Sundar S. Treatment options for visceral leishmaniasis: a systematic review of clinical studies done in India, 1980-2004. The Lancet Infectious diseases. 2005;5(12):763–74.

55. Perry MR, Wyllie S, Prajapati VK, Feldmann J, Sundar S, Boelaert M, et al. Visceral leishmaniasis and arsenic: an ancient poison contributing to antimonial treatment failure in the Indian subcontinent? PLoS neglected tropical diseases. 2011;5(9):e1227.

56. Rais S, Perianin A, Lenoir M, Sadak A, Rivollet D, Paul M, et al. Sodium stibogluconate (Pentostam) potentiates oxidant production in murine visceral leishmaniasis and in human blood. Antimicrobial agents and chemotherapy. 2000;44(9):2406–10.

57. Moreira VR, de Jesus LCL, Soares RP, Silva LDM, Pinto BAS, Melo MN, et al. Meglumine Antimoniate (Glucantime) Causes Oxidative Stress-Derived DNA Damage in BALB/c Mice Infected by Leishmania (Leishmania) infantum. Antimicrobial agents and chemotherapy. 2017;61(6).

58. Ghosh AK, Sardar AH, Mandal A, Saini S, Abhishek K, Kumar A, et al. Metabolic reconfiguration of the central glucose metabolism: a crucial strategy of Leishmania donovani for its survival during oxidative stress. FASEB journal : official publication of the Federation of American Societies for Experimental Biology. 2015;29(5):2081–98.

59. Finzi JK, Chiavegatto CW, Corat KF, Lopez JA, Cabrera OG, Mielniczki-Pereira AA, et al. Trypanosoma cruzi response to the oxidative stress generated by hydrogen peroxide. Molecular and biochemical parasitology. 2004;133(1):37–43.

60. Maldonado E, Rojas DA, Morales S, Miralles V, Solari A. Dual and Opposite Roles of Reactive Oxygen Species (ROS) in Chagas Disease: Beneficial on the Pathogen and Harmful on the Host. Oxid Med Cell Longev. 2020;2020:8867701.

61. de Oliveira J, Denadai MB, Costa DL. Crosstalk between Heme Oxygenase-1 and Iron Metabolism in Macrophages: Implications for the Modulation of Inflammation and Immunity. Antioxidants (Basel). 2022;11(5).

62. Flannery AR, Renberg RL, Andrews NW. Pathways of iron acquisition and utilization in Leishmania. Current opinion in microbiology. 2013;16(6):716–21.

63. Oexle H, Gnaiger E, Weiss G. Iron-dependent changes in cellular energy metabolism: influence on citric acid cycle and oxidative phosphorylation. Biochimica et biophysica acta. 1999;1413(3):99–107.

64. Sukhbaatar N, Weichhart T. Iron Regulation: Macrophages in Control. Pharmaceuticals (Basel). 2018;11(4).

65. Ni S, Yuan Y, Kuang Y, Li X. Iron Metabolism and Immune Regulation. Frontiers in immunology. 2022;13:816282.

66. Goto Y, Ito T, Ghosh S, Mukherjee B. Access and utilization of host-derived iron by Leishmania parasites. J Biochem. 2023;175(1):17–24.

67. Flannery AR, Huynh C, Mittra B, Mortara RA, Andrews NW. LFR1 ferric iron reductase of Leishmania amazonensis is essential for the generation of infective parasite forms. The Journal of biological chemistry. 2011;286(26):23266–79.

68. Lim D, Kim KS, Jeong JH, Marques O, Kim HJ, Song M, et al. The hepcidin-ferroportin axis controls the iron content of Salmonella-containing vacuoles in macrophages. Nature communications. 2018;9(1):2091.

69. Lafuse WP, Story R, Mahylis J, Gupta G, Varikuti S, Steinkamp H, et al. Leishmania donovani infection induces anemia in hamsters by differentially altering erythropoiesis in bone marrow and spleen. PloS one. 2013;8(3):e59509.

70. Pippard MJ, Moir D, Weatherall DJ, Lenicker HM. Mechanism of anaemia in resistant visceral leishmaniasis. Annals of tropical medicine and parasitology. 1986;80(3):317–23.

71. Londino JD, Gulick D, Isenberg JS, Mallampalli RK. Cleavage of Signal Regulatory Protein alpha (SIRPalpha) Enhances Inflammatory Signaling. The Journal of biological chemistry. 2015;290(52):31113–25.

